# TNF⍰-mediated myeloid-instructed CD14^+^CD4^+^ T cells within the tumor microenvironment are associated with poor survival in non-small cell lung cancer

**DOI:** 10.1101/2024.08.02.606259

**Authors:** Claire Marceaux, Ilariya Tarasova, Daniel Batey, Kenta Yokote, Velimir Gayevskiy, Reema Jain, Marina Leiwe, Laurie Choux, Lucy Riley, Nina Tubau Ribera, Jack Hywood, Michael Christie, Phillip Antippa, Terence P Speed, Kelly L. Rogers, Belinda Phipson, Marie-Liesse Asselin-Labat

## Abstract

The tumor microenvironment (TME) harbors a diverse array of innate and adaptive immune cells that can either support anti-tumor immunity or facilitate tumor progression. Among these, myeloid cells are key drivers of immunosuppression, yet therapeutic strategies targeting their function have achieved limited clinical success. A deeper understanding of the mechanisms by which immunosuppressive myeloid cells promote tumor progression is needed to guide the development of more effective treatments. Using spatial multi-omics analyses, we identified a population of myeloid-instructed CD14^+^ T cells present in both the tumor core and adjacent non-malignant lung tissue of patients with non-small cell lung cancer (NSCLC). CD4^+^ T cells acquired CD14 from myeloid cells by trogocytosis, resulting in an atypical T cell phenotype. High infiltration of CD14^+^CD4^+^ T cells in the tumor core was associated with poor patient survival. Spatial transcriptomics profiling revealed that tumors enriched in CD14^+^CD4^+^ T cells exhibited increased TNF⍰ signaling. Functional assays demonstrated that TNF⍰ enhanced trogocytosis, promoting the transfer of myeloid membrane components to CD4□ T cells and driving the accumulation of CD14□CD4□ T cells. These findings uncover a novel TNF⍰-mediated mechanism of immunosuppression in the TME, whereby TNF⍰ promotes aberrant myeloid–T cell interactions that may contribute to tumor progression. This work highlights a previously unrecognized axis of innate–adaptive immune crosstalk in NSCLC and suggests that targeting TNF⍰ could disrupt this pathway, restore effective T cell function, and improve therapeutic outcomes.

## Introduction

Histopathological analysis of cancer specimens is an essential part of clinical diagnosis to inform therapeutic decisions. Tissue architecture and cellular organization provide a layer of information critical for pathologists to make informed diagnoses. Similarly, the characterization of immune cell phenotypes and functional states in the physical context of a tumor is necessary to gain a broad understanding of cancer immunology. This knowledge is essential in non-small cell lung cancer (NSCLC) where immunotherapy has improved patient outcomes as single agent or in combination with chemotherapy. Yet, a number of patients do not respond to these treatments^1,2^. A comprehensive mapping of the complexity and heterogeneity of the cellular interactions in the tumor microenvironment (TME) driving tumor progression and relapse is still lacking, hindering the development of therapies tailored to the specific characteristics of individual tumor ecosystems.

Tumor cell molecular characteristics such as tumor mutational burden (TMB) and PDL1 expression are still heavily relied upon to determine patient prognostic and therapeutic approaches^3,4^. However, recent studies demonstrate the importance of assessing the immune contexture to understand tumor evolution^5,6^ and for prognostic and therapeutic purposes^7,8^. Evaluation of immune cell composition and activity in NSCLC have been performed in small patient cohorts using single-cell transcriptomics combined with cell surface protein analyses^9–12^. Detailed evaluation of individual tumor cells and immune cells determined that the intersection between tumor cell-intrinsic factors, such as *TP53* mutation and TMB, combined with immune cell characteristics may be used to better predict response to immunotherapeutics^12^. However, these studies lack spatial resolution and information on how immune cell communities may influence immune and tumor cell behaviors. Recent work by Sorin *et al.* utilized imaging mass cytometry to profile the TME in LUAD and revealed that specific immune cell neighborhoods composed of B cells and T helper cells were associated with better patient outcomes^13^. Further analyses by this group to compare LUAD and LUSC spatial architecture demonstrated the interactions between myeloid and lymphoid cells as a prominent feature of LUAD^14^. These studies highlight how cellular communities and their specific location, identified through spatial analysis, have a stronger biological impact than individual cells. Evaluating this spatial organization provides a new layer of information, akin to that utilized in anatomical pathology for diagnosis, to decipher tumor biology.

Tumor-infiltrating myeloid-derived monocytes or neutrophils, called myeloid-derived suppressor cells, and tumor-associated macrophages are key drivers of immunosuppression in cancer^15,16^. While the role of myeloid cells in blocking anti-tumor immunity has been recognized for a long time, there has been limited success in large-scale clinical trials of therapies that modulate myeloid cell function^17^. Preclinical and translational studies in organ-specific tumor contexts are essential in understanding the many mechanisms by which myeloid cells drive immunosuppression in the TME, and exploiting myeloid cell vulnerabilities to increase the success of myeloid-targeted therapies. Recent studies have highlighted that myeloid cell-mediated immunosuppression can be initiated in early myeloid cell progenitor cells in the bone marrow^18^, as well as locally, within the TME, through the instruction of lymphoid cells^18^. LaMarche *et al.*^19^ showed that basophil-derived IL-4 reprogrammed myeloid progenitor cells in the bone marrow of NSCLC tumor-bearing mice, supporting the differentiation of tumor-promoting myeloid cells. Depleting IL-4 signaling in myeloid progenitor cells in the bone marrow using conditional knockout mice reduced tumor burden^19^. Systemic inhibition of IL-4 with Dupilumab in combination with immune checkpoint inhibitors reduced circulating monocytes and increased CD8^+^ T cell infiltration in a small clinical trial^19^ demonstrating the effect of tumor cells beyond the local TME.

The impact of myeloid cells in modulating local adaptive immunity was evaluated by Pallett *et al*.^18^ where they described a new subset of myeloid-instructed tissue-resident T cells present in the skin, lymph nodes, and liver. They showed that a subset of CD8^+^, and in lower abundance CD4^+^ T cells, expressed the TLR4 coreceptor CD14, a canonical marker of myeloid cells and innate immunity^18^. These myeloid-instructed T cells were reprogrammed by the transfer of myeloid material to T cells upon direct contact between T cells and mononuclear phagocytes and were found to accumulate in hepatitis B virus-infected livers and in hepatocarcinomas. The existence and contribution of these myeloid-instructed T cells to anti-tumor immunity in solid tumors is still unknown. In addition, whether tumor signaling to the bone marrow could promote myeloid progenitor cells to differentiate into an ‘instructor’ phenotype, or whether this is mediated by local tissue-specific signal remains to be elucidated.

Here we applied multiplex ion beam imaging (MIBIscope) and spatial transcriptomic (GeoMx) to a clinically annotated cohort of NSCLC patient samples to identify cellular and molecular events controlling anti-tumor immunity. We demonstrated the existence of a subset of clonally expanded T cells expressing the myeloid marker CD14. Analysis of three distinct cohorts of NSCLC confirmed the existence of this discrete T cell subset in tumors, and showed that high infiltration of CD14^+^CD4^+^ T cells in the tumor core correlated with short-term patient survival. CD4^+^ T cells acquired CD14 from myeloid cells by trogocytosis, resulting in an atypical T cell phenotype. Tumors enriched in myeloid-instructed CD14^+^CD4^+^T cells exhibited an upregulation of genes in the TNF⍰ signaling pathway. We showed that TNF⍰ increased CD4^+^ T cell trogocytosis, augmenting the proportions of myeloid-instructed T cells. TNF⍰ is known to exert a number of pro-tumorigenic activities^20,21^. Here, we show that TNF⍰-induced trogocytosis and myeloid instruction of T cells may be an additional mechanism by which TNF⍰ reduces anti-tumor immunity.

## Results

### A longitudinal cohort of resected NSCLC with survival outcome

To study the features of anti-tumor immunity associated with patient outcome in resectable NSCLC, we utilized a bank of 136 treatment-naïve NSCLC samples (stage I to IIIa) for which we had up to 10 years of longitudinal clinical follow-up (Fig 1A). To characterize the cellular traits of the TME that may be associated with short or prolonged survival, we selected tissue samples from patients with an overall survival of less than 3 years or more than 6 years post-surgery to perform spatial proteomics and spatial transcriptomics analyses. A total of 68 patients were analyzed, comprising 28 samples with less than 3 years survival (21 LUAD and 7 LUSC) and 40 samples (26 LUAD and 14 LUSC) with more than 6 years survival (Figure 1A). H&E images were annotated by a pathologist to select regions of interest (ROIs) for spatial single-cell proteomics using the MIBIscope and spatial transcriptomics using whole transcriptome profiling on the GeoMx. On each ROI, GeoMx enabled the analysis of the gene expression profile of tumor cells exclusively (area of illumination (AOI) panCK^+^ cells) and the immune compartment (AOI CD45^+^ cells) (Fig 1B). After quality control checks, a total of 920 field-of-views (FOVs: 400×400μm, average of 5 per patient) MIBI scans and 234 matched AOIs (average of 4 per patient) on GeoMx were selected for further analysis corresponding to 64 patients (44 LUAD and 20 LUSC; Fig 1A, Supp Table 1 and 2).

**Figure 1:**
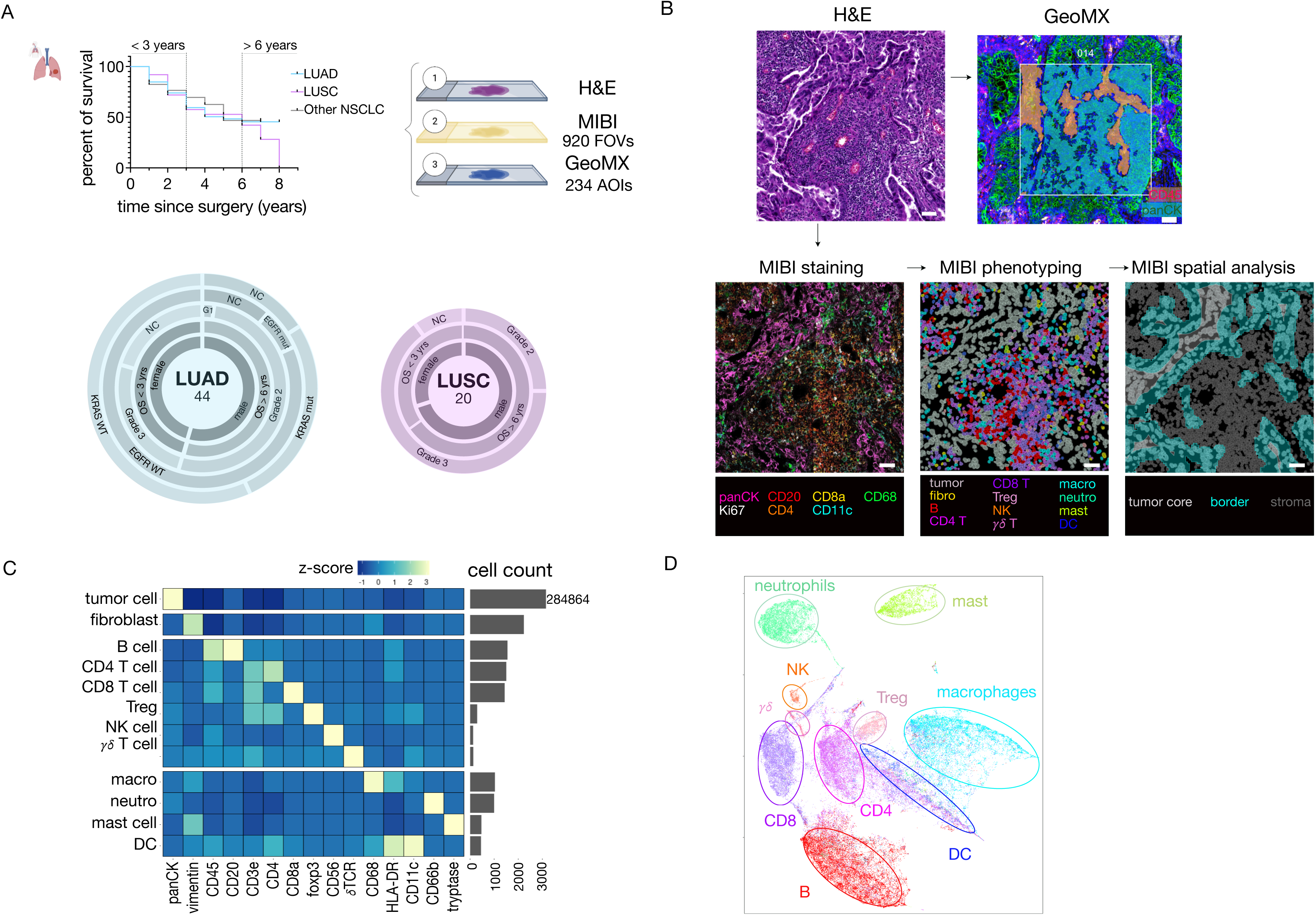
Spatial analysis of the tumor microenvironment and tumor cells in human non-small cell lung cancer samples. **A.** Top: Kaplan Meier survival curve showing overall survival of resected non-small cell lung cancer (NSCLC) sample cohort. Samples from patients with overall survival (OS) less than 3 years or more than 6 years were selected for this study. Bottom: Donuts graph representing the characteristics of the lung adenocarcinoma (LUAD) and squamous cell carcinoma (LUSC) patients analyzed. The numbers represent the final sample size of patients profiled. Layers are unrelated. G: grade; NC: not cited. **B.** Overview of the workflow for the spatial analysis. Regions of interest (ROI) were selected by a pathologist based on H&E stain. Adjacent slides were used for spatial proteomic (MIBI) and spatial transcriptomic (Nanostring GeoMx). For GeoMx analysis, selection of areas of illumination (AOI) was based on the protein expression of CD45 (pink) and cytokeratin (panCK, green). Scale bar: 50μm. For MIBI scanning, coordinates of a selected ROI were transposed to the antibody-stained slide and scanned. Scale bar: 50μm. **C.** Heatmap depicting the mean expression of each marker (or column) z-score normalized across all the cell types (rows) identified by the XGBoost model. Right panel shows the number of cells in each category. **D.** UMAP of immune cell types generated in all samples using the mean intensity measurement of each phenotypic marker in all immune cells. The colors reflect the phenotype assigned by the XGBoost model.

For spatial proteomic analysis, a bespoke panel of 38 conjugated antibodies was developed to identify tumor, immune cells and fibroblasts and detail their functional state (Supp Fig 1). For each patient, FOVs were scanned and stitched together to enable spatial analysis of cell communities in different tumor compartments: tumor core, tumor border and surrounding stroma, classified based on a pixel classifier trained in QuPath^22^ (Fig 1B). Cells were segmented using a Deep Cell model^23^ trained based on the expression of cytoplasmic (Major histocompatibility Complex Class I (MHC-I)), cell membrane (pan-cytokeratin (panCK)) and nuclei (dsDNA) markers. Individual cell phenotypes were manually annotated on 116 FOVs to train an XGboost machine learning method to phenotype twelve main cell types with an overall accuracy of 90%. A second independent model was developed for determining positive or negative expression of individual functional markers for each cell type. Using this approach, a total of 384,808 cells were classified into epithelial cells, fibroblasts and ten major immune cell subpopulations: B cells, CD4^+^ conventional T cells, CD8^+^ T cells, CD4^+^ regulatory T cells, natural killer cells, gamma-delta T cells, macrophages, neutrophils, mast cells and dendritic cells. A heatmap of the mean expression of markers used for cell type annotation indicated a strong correlation between cell classification and the expression of expected markers (Fig 1C). Classified cells corresponded to 73% tumor cells, 21% immune cells and 6% stromal cells. Unsupervised clustering of immune cells demonstrated the performance of our classification model with all lymphoid cells clustering away from myeloid cell subsets (Fig 1D). Consistent with previous publications utilizing *in situ* methods or tissue dissociation approaches^5,11,13,24^, the five most abundant immune cell types infiltrating NSCLC samples were B cells, CD4^+^ conventional T cells, CD8^+^ T cells, macrophages and neutrophils (Fig 1C).

Overall we have exploited a cohort of clinically annotated NSCLC samples to develop methods for tissue classification and cell-type phenotyping from single-cell multiplex protein imaging, and to interrogate the association between spatial proteomics and transcriptomics data with clinical outcomes.

### Distinct spatial clusters of immune cells in the TME of LUAD and LUSC

We first investigated the characteristics of the tumor cells in LUAD and LUSC tissues. Consistent with previous studies^25,26^, GeoMX analysis of panCK^+^ tumor cell regions showed clear transcriptomic differences between LUAD and LUSC tumor cells as well as heterogeneity in the gene expression profile of tumor cells between patients of the same histological subtypes (Fig 2A). LUAD and LUSC expressed typical phenotypic markers of each histological subtypes (eg. *NKX2.1*, *KRT7*, *MUC1*, *SFTPB* in LUAD and *KRT5*, *KRT6*, *TP63* in LUSC; Supp Fig 2A). With the proteomics analysis, we refined the co-expression of distinct functional markers in individual tumor cells. We observed a heterogenous pattern of expression of MHC-I and MHC-II in the two cancer subtypes (Fig 2B), consistent with previous studies where ∼50% of tumors were found to have lost or had low levels of MHC-I and MHC-II^27,28^. MHC-I and MHC-II expressions were positively correlated in LUAD (Spearman correlation score: 0.7). Interestingly, loss of MHC-I was often associated with reduced expression of IFNγ (Spearman correlation score: 0.49) in LUAD tumor cells, but not in LUSC (Fig 2B). In LUSC, MHC-II expression was negatively associated with Ki67 expression (Spearman correlation score: -0.49). Expression of PDL1 in tumor cells varied across patients but was consistently lost in MHC-I negative tumors (Spearman correlation score: 0.32 in LUAD). IFNγ expression was low in all LUSC samples, except for one patient, whereas ∼50% of LUAD tumors expressed IFNγ (Fig 2B). In LUAD, MHC-I/II^low^ tumors had less immune cell infiltration in the tumor core or border than MHC-I/II^high^ tumors. The tumor cell-specific expression of each of these markers, including PDL-1, was not associated with relapse-free survival or overall survival (Fig 2B and Supp Fig 2B).

**Figure 2.**
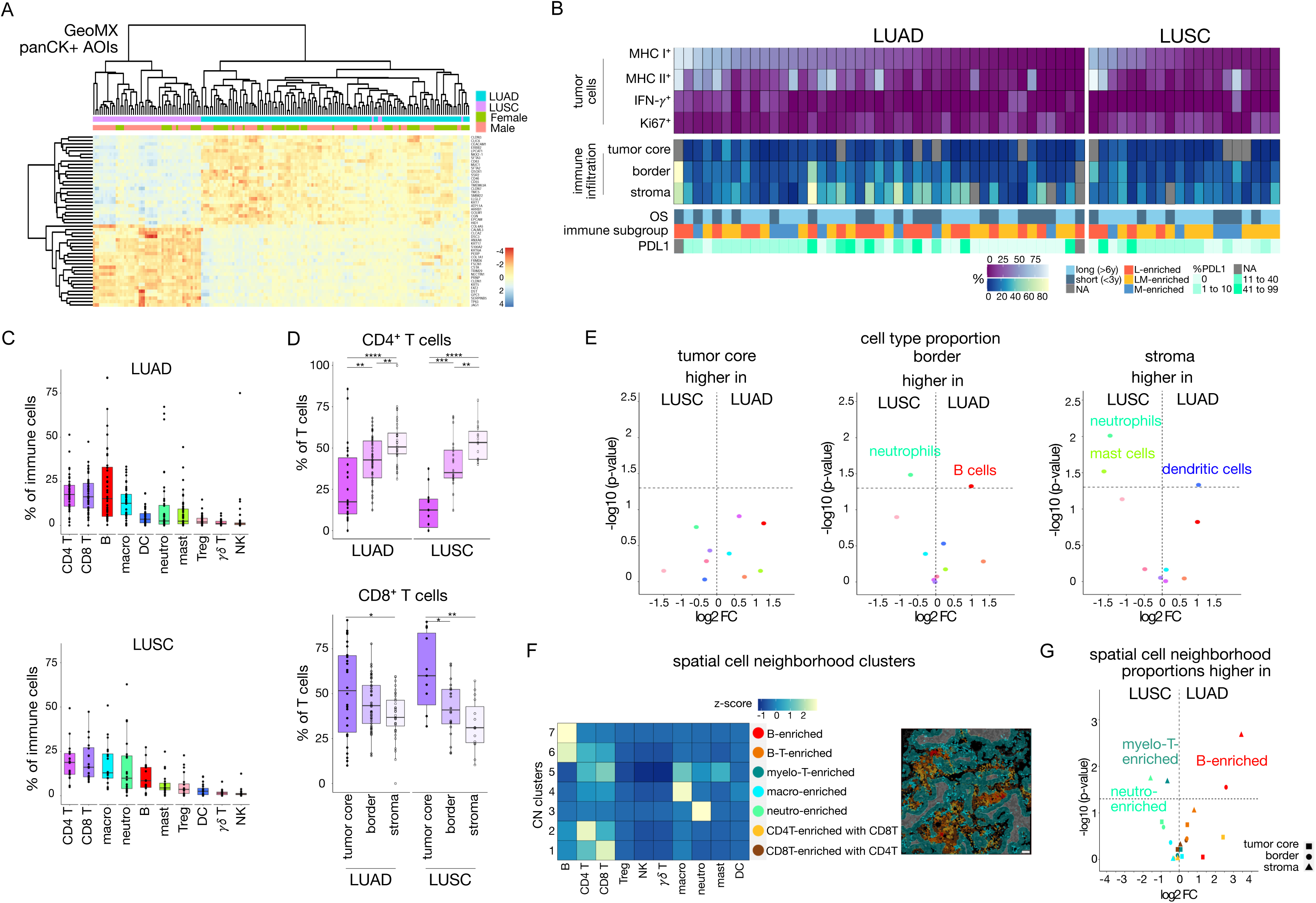
Cell communities containing myeloid cells are a feature of the LUSC TME. **A.** Heatmap showing the expression of the top 50 differentially expressed genes in cytokeratin positive tumor cells (panCK^+^ AOI, GeoMx) between LUAD and LUSC. Data show z-scores. **B.** Top: Heatmap depicting the proportion of MHC I^+^, MHC II^+^, IFNγ^+^, and Ki67^+^ in tumor cells (PanCK^+^) in individual patient samples from MIBI analysis (percentages). Samples ordered highest to lowest relative to the proportion of MHC I^+^ cells. Middle: heatmap representing the proportion of immune cell infiltration in the three tissue compartments (tumor core, border, stroma) in individual patients (percentages). Bottom: Clinical and molecular data on individual patients. OS: overall survival. **C.** Proportion of the different immune cell types in LUAD and LUSC, as determined by MIBI. Data show median ± SD. n=44 LUAD, n=20 LUSC. **D.** Boxplots showing the localization of CD4^+^ T cells and CD8^+^ T cells in the different spatial tissue compartments in LUAD and LUSC patients. Data show median ± SD. ****, p<0.0001; ***, p<0.001; **, p<0.01; *, p<0.05 comparing the different compartments, Wilcoxon test. n=44 LUAD, n=20 LUSC. **E.** Volcano plots comparing the proportion of each immune cell subtype in the three compartments in LUAD and LUSC samples. A horizontal dashed line indicates -log10(0.05), Wilcoxon test. n=44 LUAD, n=20 LUSC. **F.** Heatmap showing seven cell neighborhoods (CN) detected in NSCLC samples, and the cell type composition for each cluster. Right panel: representative images showing rendering of different CN in tumor core (grey), border (blue) and stroma (dark grey). Scale bar: 50μm. **G.** Volcano plots comparing the proportion of each CN cluster in the three compartments between LUAD and LUSC samples. A horizontal dashed line indicates -log10(0.05), Wilcoxon test.

We then compared the immune microenvironment of LUAD and LUSC samples. In both tumor subtypes, the predominant immune subsets in the whole tissue were CD4^+^ and CD8^+^ T cells, followed by B cells in LUAD, and macrophages in LUSC (Fig 2C). Analysis of the specific location of the cells in different tumor compartments – tumor core, tumor border or stroma – showed that immune cells were more frequent in the stroma (Supp Fig 2C). Within the immune cells, the proportions of distinct immune subsets were evenly distributed between the compartments or enriched in the stroma, except for macrophages and CD8^+^ T cells that were more frequently detected in the tumor core and at the tumor border, in both LUAD and LUSC (Fig 2D and Supp Fig 2D). Whole tissue analysis of immune cell infiltration revealed significantly higher B cell infiltration in the LUAD TME compared with LUSC, whereas neutrophils were more often present in LUSC (Supp Fig 2E). Detailed analysis of the TME in different compartments of the tumors highlighted that the enrichment for B cells in LUAD occurred at the border of the tumor front, while neutrophils were located at the border and in the stroma of LUSC (Fig 2E). In addition, this analysis of the different tumor compartments highlighted further differences between the two histological subtypes where dendritic cells were found enriched in LUAD stroma compared with LUSC and mast cells were mostly present in the LUSC stroma compared with LUAD (Fig 2E). Together, these results point to a more myeloid-enriched microenvironment in LUSC compared with LUAD.

Analysis of immune cell neighborhoods (CN) revealed 7 spatial clusters based on their immune composition, with neighborhoods of a single cell type (CN7: B cell, CN4: macrophages and CN3: neutrophils), T cells (CN1 and 2 with higher prevalence of CD8 or CD4 respectively), lymphoid cells (CN6: B and T cells) and a mix of myeloid and T cells (CN5: T cells with macrophages and mast cells) (Fig 2F). Evaluating the presence of these clusters in LUAD and LUSC, we observed B cell-enriched CN7 in the stroma and at the border of LUAD, whereas neutrophil-enriched CN3 was present in the stroma of LUSC (Fig 2G), indicating that these enriched cell populations observed in each tumor subtypes infiltrate the tumor as cell communities and not as individual cells. Interestingly the myeloid/T cell cluster was enriched in the stroma of LUSC compared with LUAD (Fig 2G).

Together, these analyses revealed key differences in the spatial organization and composition of the TME in LUSC and LUAD, highlighting essential immune cell communities characteristic of each tumor type.

### Stratification of NSCLC according to myeloid and/or lymphoid cell infiltration is not associated with oncogenic driver mutations

We then investigated the heterogeneity of the TME between individual tumors in LUAD and LUSC samples. Quantification of the proportions of lymphoid and myeloid cells in each sample followed by unsupervised clustering of samples resulted in the stratification of patients into 2 subgroups in LUSC (Supp Fig 3A) and 3 subgroups in LUAD (Fig 3A, B). In LUSC, 15 tumors were characterized by a mix of lymphoid and myeloid cell infiltration (LM-enriched, 75%), with a TME rich in T and B cells with macrophages, mast cells and dendritic cells. Five LUSC tumors were classified as myeloid-enriched with little lymphoid cell infiltration (M-enriched, 25%) (Supp Fig. 3A). In LUAD, most tumors had high lymphoid infiltration (79.5%) either alone (L-enriched, n=19, 43%) or mixed with myeloid cell infiltration (LM-enriched, n=16, 36%) (Fig 3A). Myeloid-enriched tumors only represented 20.5% of the LUAD samples (M-enriched, n=9). We then analyzed two independent cohorts of NSCLC samples in which the tumor immune infiltrate was evaluated using Image Mass Cytometry^13,29^, a hi-plex mass-spectrometry-based imaging approach similar to the one we employed with the MIBIscope. Sorin *et al.* analyzed 416 LUAD samples with a 35-plex antibody panel to characterize cancer cells, stromal cells and innate and adaptive immune cells^13^, while Enfield *et al*. analyzed 39 LUAD and 23 LUSC lesions with two distinct antibody panels to interrogate these cell types^29^. Consistent with our cohort, unsupervised clustering based on proportions of distinct immune cell infiltration led to the stratification of LUAD tumors in L-, LM- and M-enriched tumors (Fig 3A and B). In the two cohorts where genomic profiling was available (Marceaux *et al*, and Enfield *et al*.), *EGFR^mut^* or *KRAS^mut^* tumors were found in each subgroup: 2 *EGFR^L858R^* (5.7%) and 8 *KRAS^mut^* (22.9%) in the 35 L-enriched tumors, 2 *EGFR^L858R^* (6.5%) and 9 *KRAS^mut^* (29%) in the 31 LM-enriched, 2 *EGFR^L858R^*(11.8%) and 7 *KRAS^mut^* (41.2%) in the 17 M-enriched tumors (Fig. 3B). *STK11^mut^* was only observed in one tumor in the L-enriched group. These results indicate that the type of immune infiltration in LUAD is not solely dictated by the presence of oncogenic driver mutations.

**Figure 3.**
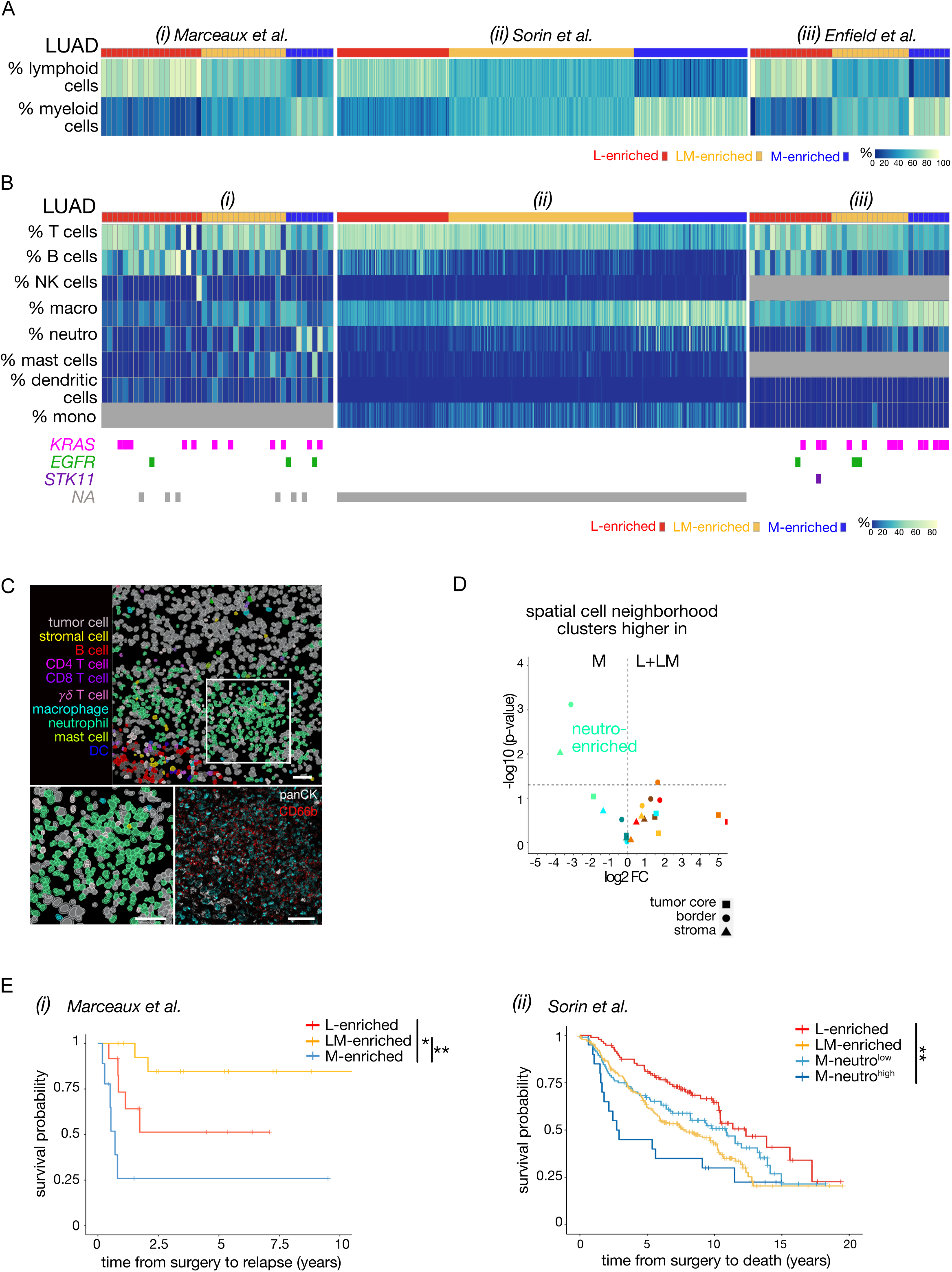
Enrichment for neutrophil clusters is associated with worst relapse-free survival in LUAD patients. **A and B. (A)** Heatmaps showing the proportion of lymphoid and myeloid cells in all immune cells in individual samples in LUAD in our cohort (i), in Sorin *et al*. cohort (ii) and in Enfield *et al*. cohort (iii). K-mean clustering stratified the TME in LUAD patients in the 3 cohorts in 3 immune subgroups (k=3) lymphoid-enriched (L-enriched; n=19 in (i); n=113 in (ii); n=16 in (iii)), lymphoid/myeloid-enriched (LM-enriched; n=16 (i), n=118 (ii), n=15 (iii)) or myeloid-enriched (M-enriched; n=9 (i), n=115 (ii), n=8 (iii)). (**B**) Heatmaps showing the proportions of immune subpopulations for each patient in the 3 different immune subgroups. *KRAS*, *EGFR* and *STK11* mutations are specified for each sample. ‘NA’: not available. **C.** Representative image showing clustering of neutrophils in a LUAD tumor (MH1041). Scale bar: 50μm. **D.** Volcano plots comparing the proportion of each CN cluster in the three compartments between M- and L+LM-enriched LUAD samples. A horizontal dashed line indicates -log10(0.05), Wilcoxon test. **E.** Kaplan-Meier survival curve showing (i) relapse-free survival in the three groups of our LUAD patients (*, p<0.05, **, p<0.01 Cox proportional hazards regression model (Wald test). L-enriched, n=19; LM-enriched, n=16; M-enriched, n=9) and (ii) overall survival in the immune subgroups defined previously using Sorin *et al.* where M-enriched patients have been split into low (M-Neutro^low^) and rich (M-Neutro^high^) neutrophil environment (**, p<0.01 Cox proportional hazards regression model (Wald test). L-enriched, n=113; LM-enriched, n=118; M-Neutro^low^-enriched, n=95; M-Neutro^high^-enriched, n=20).

### Neutrophil infiltration is associated with worse patient outcomes in LUAD

We then assessed differences between LUAD immune subgroups where we had the largest number of samples. L-enriched tumors had high T and B cell infiltration, although high B or T cell infiltration appeared to be mostly mutually exclusive (Fig. 3B). LM-enriched tumors had lower B-cell infiltration than L-enriched tumors and increased infiltration of macrophages (Fig. 3B). We combined all patients with tumors harboring lymphoid infiltration (L-enriched and LM-enriched) as the LLM-enriched subgroup for further analysis. Only when quantifying immune cell infiltration in the individual tissue compartments did we observe that tumor cores infiltrated with B cells were significantly associated with long RFS (Supp Fig 3B), consistent with previous work describing B cells infiltration as a predictor of long survival^13^.

In M-enriched tumors, the dominant populations were macrophages and neutrophils. Each myeloid cell subtype was located equally in the different tumor compartments with the exception of dendritic cells that were more often found at the tumor border or in the stroma in LUAD (Supp Fig 3C). The expression of functional markers of myeloid cells in LUAD was similar in each individual compartment (Supp Fig 3D). A distinct feature of M-enriched tumors compared with LM-enriched tumors was a higher infiltration of neutrophils. Cell neighborhood analysis revealed that in all compartments of M-enriched tumors, neutrophils were present in clusters compared with L- and LM-enriched tumors (Fig 3C, D and Supp Fig 3E). In our cohort, five out of the nine patients with an M-enriched LUAD had high neutrophils infiltration and an RFS of less than 7 months, significantly shorter than patients with L and LM-enriched LUAD (median RFS 20.3 months and 64.9 months, respectively, Fig 3E, Supp Fig. 3F). Of note, the long survivor with an M-enriched tumor had an *EGFR^L858R^* mutation characterized by a distinct TME compared with the other M-enriched tumors, with enrichment in mast cells and low levels of neutrophil infiltration (Supp Fig 3G). This observation is consistent with previous results showing that increased mast cells in the TME is associated with never-smoker patients and prolonged survival^13^. In Sorin *et al*.^13^, patients with M-enriched, high neutrophil infiltrated tumors were found to have worse overall survival consistent with our cohort and previous studies^29^ (Fig 3E). Neutrophil-enriched tumors also had worse survival in LUSC (Supp Fig. 3H).

Overall, these results demonstrate the inter-tumor heterogeneity in the TME of NSCLC, enabling the stratification of patients according to different types of immune infiltrate. We identified a subset of M-enriched tumors with a TME rich in neutrophils that was not observed when lymphoid cells were also present in the TME. Patients with these distinct neutrophil-enriched tumors had worse clinical outcomes, whereas LLM-enriched tumor with B cell infiltration specifically in the tumor core had better RFS.

### CD14^+^CD4^+^ and CD14^+^CD8^+^ T cells are present in the microenvironment of LUAD and LUSC tumors

With the recent observation that myeloid cells can instruct T cells in hepatocarcinoma through the transfer of membrane material^18^, we evaluated whether CD14^+^ T cells were also a feature of NSCLC. Quantification of CD14 expression in immune cell subsets in LUAD and LUSC tumors revealed that a subset of T cells expressed CD14 (Fig 4A, B, Supp Fig 4A, Supp Fig 4B), the co-receptor for TLR-4–mediated LPS recognition usually expressed by myeloid cells, indicating that these cells may be present in more organs than the liver, skin and lymph node as described previously^18^. CD14^+^ T cells were also detected in the Sorin and Enfield datasets (Supp Fig. 4C, data not shown). In LUAD and LUSC, CD14^+^CD4^+^ T cells were present at a higher frequency than CD14^+^CD8^+^ T cells (Fig 4C, Supp Fig 4D), contrary to the results described in the liver where CD14^+^CD8^+^ T cells were predominant^18^. CD14^+^ lymphoid cells represented 8.5% to 15% of CD8^+^, Treg and CD4^+^ T cells in LUAD (Fig 4C). As a proportion of their parent population, CD14^+^CD4^+^ T cells were present equally in the tumor core, at the border or stroma (Supp Fig 4E). The presence of CD14^+^ T cells in NSCLC was further confirmed by flow cytometry analysis of fresh human lung tumors (Fig 4D and Supp Fig 4F). Image cytometry and Hoechst 33342 analysis confirmed that CD14^+^CD4^+^ T cells were single cells and not a complex of monocytes/T cells as detected in the blood^30,31^ (Fig 4E and Supp Fig 4G).

**Figure 4.**
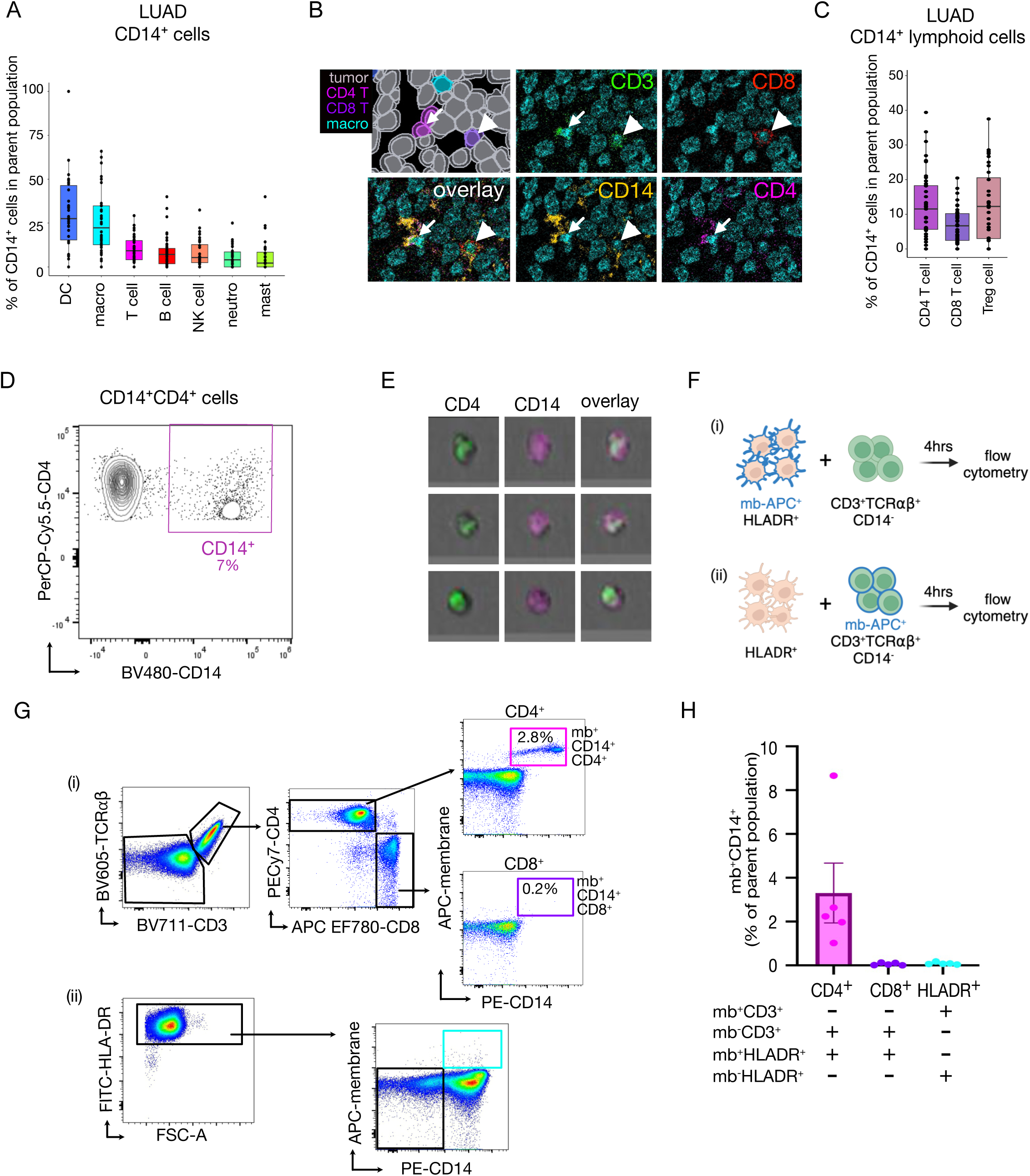
Myeloid-instructed CD14^+^ T cells infiltrate tumors in non-small cell lung cancer. **A.** Proportion of immune cell types expressing CD14 in LUAD as determined by MIBI. Data show median ± SD. n=44 LUAD. **B.** Representative MIBI images showing the expression of CD14, CD3, CD8 and CD4 in a representative LUAD tumor (AH0353). Arrow shows CD14 expression in CD3^+^CD4^+^ cells. Arrowhead shows CD14 expression in CD3^+^CD8^+^ T cells. **C.** Box plot showing the proportion cells expressing CD14 in lymphoid cells in LUAD as determined by MIBI. Data show median ± SD. n=42 LUAD. **D.** Representative flow cytometry plot showing CD14 expression in CD4^+^ T cells gated on CD45^+^CD3^+^ cells in a LUAD sample (MH0069). **E.** Representative image cytometry images showing the expression of CD14 and CD4 on CD45^+^CD3^+^CD4^+^ cells. n=2. **F.** Schematic representation of trogocytosis assay using PBMC: (i) HLADR^+^ cells were labelled with membrane-APC dye and co-cultured 4hrs with CD3^+^TCR⍰β ^+^CD14^-^ cells or (ii) HLADR^+^ cells were co-cultured 4hrs with CD3^+^TCR⍰β ^+^CD14^-^ cells labelled with membrane-APC dye, followed by flow cytometry to evaluate membrane transfer. **G**. Representative flow cytometry plots showing in (i) the APC membrane transfer from HLADR^+^ cells to CD4^+^ T cells only and (ii) the non-transfer of membrane from T cells, showing no expression of APC by HLADR^+^ cells. **H**. Quantification of the flow cytometry trogocytosis experiment showing the percentage of membrane^+^CD14^+^ in CD4 T cells, CD8 T cells and HLADR^+^ cells in the indicated conditions (n=5).

To evaluate the process by which the co-expression of a myeloid marker and T cell markers could occur, we performed trogocytosis experiments where membrane-labelled T cells (CD3^+^TCRab^+^) were incubated with HLA-DR^+^ cells or vice-versa in which membrane-labelled HLA-DR^+^ cells were incubated with CD3^+^TCRab^+^ cells (Fig 4F). Transfer of membrane only occurred from HLA-DR^+^ cells to T cells, and was most predominant in CD4^+^ T cells (Fig 4G and H), indicating that only T cells can take up membrane from myeloid cells.

Overall, these data reveal the existence of a discrete subset of CD4^+^ T cells harboring the myeloid marker CD14 in NSCLC and that this phenomenon may be mediated by T cell trogocytosis from HLA-DR^+^ myeloid cells.

### CD14^+^ T cells are not detected in healthy lungs and are located within myeloid cell neighborhoods in LUAD

We then assessed whether CD14^+^CD4^+^ T cells were a unique feature of tumors or were also present in the non-malignant or healthy lung tissues. CD14^+^CD4^+^ T cells were observed in the non-malignant lung of lung cancer patients in proportions similar to what was observed in matched tumors (Fig 5A). However, CD14^+^CD4^+^ T cells were not detected in four lung samples obtained from healthy donors (Fig 5A), indicating that CD14^+^CD4^+^ T cells may be induced in an oncogenic context. Consistently, the vast majority of CD14^+^CD4^+^ T cells expressed CD39 (Fig 5C), a marker of tumor-specific lymphocytes^32,33^, whereas only a subset of CD14^-^CD4^+^ T cells expressed CD39, suggesting that CD14^+^CD4^+^ T cells are enriched for tumor-specific T cell clones (Fig 5B). To assess the clonality of CD14^+^ T cells, we compared the TCRβ repertoire of CD14^+^CD4^+^ and CD14^-^CD4^+^ T cells by TCR sequencing of sorted cell subsets. Sequencing results showed an expansion of some TCRβ clones in CD14^+^ cells compared to CD14^-^ cells (Supp Fig 5A), indicating CD14^+^CD4^+^ T cells are an expanded population of T cell clones present in tumors although the tumor specificity of these clones can not be concluded.

**Figure 5.**
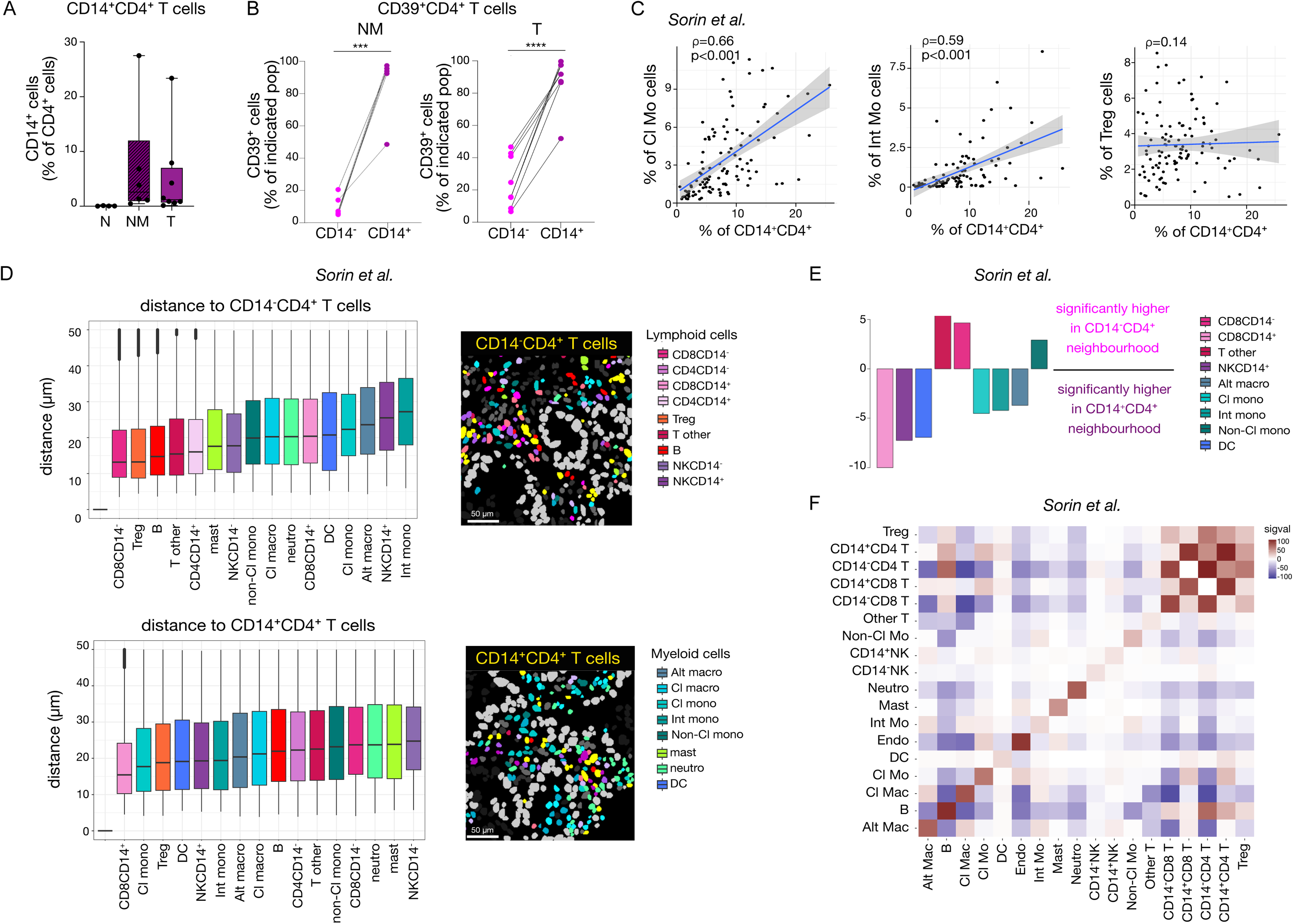
Myeloid-instructed CD14^+^CD4^+^ T cells reside in a myeloid-enriched microenvironment. **A.** Flow cytometry quantification of percentage of CD14^+^CD4^+^ T cells in normal lung (N) (n=4), non-malignant lung (NM) (n=6) and tumor (T) (n=8). **B.** Proportion of CD14^+^CD4^+^ and CD14^-^CD4^+^ T cells expressing CD39 in non-malignant lung tissues (NM) (n=5) and tumor tissues (T) (n=7). n=4 LUAD, n=3 LUSC and n=1 carcinoid. ***, p<0.001, **, p<0.01; *, p<0.05, paired t-test between CD14^+^ and CD14^-^ T cell subsets. **C.** Correlation plots between the proportion of CD14^+^CD4^+^ T cells and classical monocytes (Cl Mo), intermediate monocytes (Int Mo) and regulatory T cells (Treg) in Sorin *et al.* L-enriched immune subgroup. Spearman rank correlation. n=113 LUAD. **D.** Boxplots showing the distance between indicated cell type and CD14^-^CD4^+^ T cells (top left) or CD14^+^CD4^+^ T cells (bottom left) within 50µm radius in Sorin *et al.* L-enriched subgroup (n=112 LUAD). Right panel shows representative images of segmented cells in the microenvironment around CD14^-^CD4^+^ T cells (yellow, top) and CD14^+^CD4^+^ T cells (yellow, bottom). **E.** Output of Propeller analysis showing which cell type proportion is significantly higher in CD14^-^CD4^+^ T cells and CD14^+^CD4^+^ T cells neighborhood in Sorin *et al.* L-enriched subgroup (n=112 LUAD). The Y-axis shows t-statistics after filtering by FDR<0.05. **F.** Heatmap representative of cell-cell interaction compared to random permutations distribution in Sorin *et al.* cohort L-enriched immune subgroup (n=113 LUAD). Heatmap showing the summed significance scores for each cell type pair across all 113 images. For each IMC image, cell type pairs were assigned a score based on spatial interaction: 1 for interaction (red), 0 for no interaction or avoidance (white), and −1 for avoidance (blue). These scores were then summed across all images, resulting in a possible range from –113 to 113.

To better understand the context in which CD14^+^CD4^+^ T cells arise, we investigated whether the presence of CD14^+^CD4^+^ T cells was correlated with abundance and spatial organization of other immune cell types. For this analysis, we exploited the Sorin *et al*. dataset^13^, given the large number of samples available. Proportions of CD14^+^CD4^+^ T cells were correlated with the presence of classical and intermediate monocytes, two populations of CD14-expressing immune cells (Fig 5C), as well as the proportion of CD14^+^CD8^+^ T cells (Supp Fig 5B). However, an inverse correlation was observed with CD14^-^CD4^+^ and CD8^+^ T cells (Supp Fig 5B). The proportion of CD14^+^CD4^+^ T cells was not correlated with the presence of other lymphoid cell types, including immunosuppressive Tregs, NK or B cells (Fig 5C and Supp Fig 5B), nor with the presence of neutrophils, mast cells or dendritic cells (Supp Fig 5B).

To define the cellular niche in which CD14^+^CD4^+^ T cells may be found, we first quantified cell-cell distances to CD14**^+^**CD4^+^ T cells and CD14^-^CD4^+^ T cells within each sample. These analyses revealed that CD14^+^CD4^+^ T cells were in close proximity to myeloid cells and other CD14-expressing cells, whereas CD14^-^CD4^+^ T cells were closer to lymphoid cells (Fig 5D, E). Analysis of cell-cell interaction shows that homotypic interactions, where identical cell types interact with each other, is a clear feature of the NSCLC TME (Fig. 5F). T cells in general, including Treg, had more cell interactions between themselves than any other cell types, with the CD8^+^ and CD4^+^ T cell interactions being the strongest after homotypic interactions. CD14^-^CD4^+^ T cells interacted more with CD14^-^CD8^+^ T cells than CD14^+^ T cells, while CD14^+^CD4^+^ T cells interacted more with CD14^+^ T cells and myeloid cells than CD14^-^ T cells. Amongst their myeloid cell interactions, CD14^+^CD4^+^ T cells showed attraction towards monocytes and macrophages, but little attraction to DC and neutrophils. B cells interacted with other lymphoid subsets, but showed repulsion with myeloid cell subsets including macrophages, monocytes and neutrophils (Fig. 5F).

Overall, these spatial analyses indicate that CD14^+^CD4^+^ T cells evolve in a myeloid-enriched niche that favors interactions with other CD14-expressing T cells, macrophages and monocytes.

### CD14□CD4□ T cells represent a distinct, myeloid-instructed T cell population associated with poor prognosis and TNF⍰-driven signalling

Trogocytosis is a dynamic process that involves the transfer of plasma membrane and its associated cytosol between two cell types, resulting in changes in the functional properties of the recipient cell^34^. Functional characterization of CD14^+^CD4^+^ T cells using spatial proteomics data revealed CD14^+^CD4^+^ T cells express other prototypic markers of myeloid cells (MHC-II, CD16, CD163, CD206), and a marker of tissue residency (CD49a) (Fig. 6A) but lower expression of CD103 (Supp Fig 6B). Flow cytometry data further showed that CD14^+^CD4^+^ T cells have a distinct activation phenotype to their CD14-counterparts, exhibiting elevated expression of Ki67, IFNγ and OX40 but low expression of canonical co-activation (ICOS) and exhaustion marker (PD1) or T cell activation (CD69) (Fig. 6B and Supp Fig. 6A). While this profile does not align with classical effector or helper T cell subsets, it may reflect a tumor-adapted state associated with an inflammatory environment.

**Figure 6.**
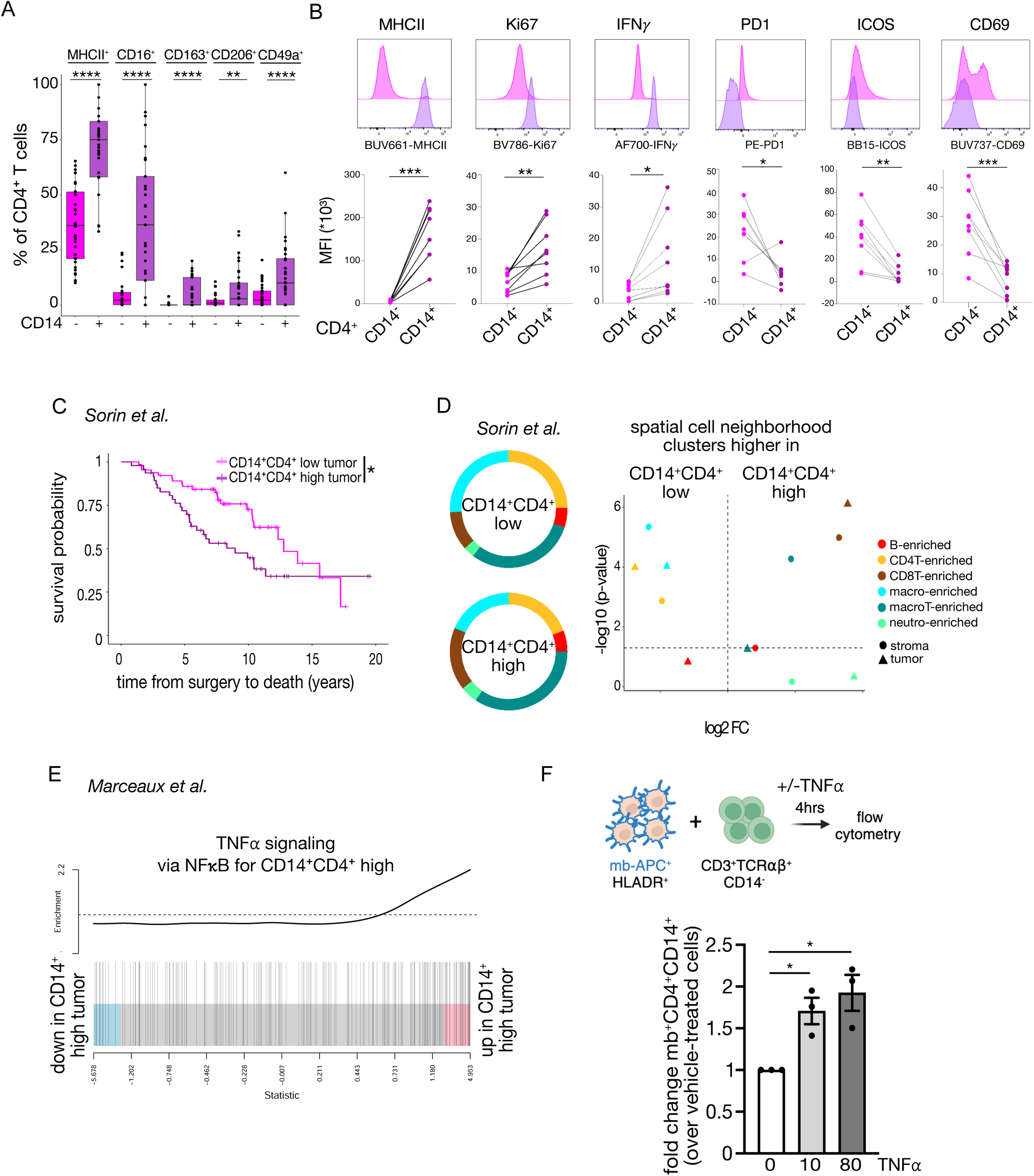
High infiltration of CD14^+^CD4^+^ T cells is associated with poor survival and high TNF⍰ signalling. **A.** Expression of MHC-II, CD16, CD163, CD206 and CD49a in CD14^+^CD4^+^, CD14^-^CD4^+^ T cells as determined by MIBI. ***, p<0.001, **, p<0.01; *, p<0.05, paired t-test. Data show median ± SD. n=42 LUAD. **B.** Mean fluorescence intensity (MFI) of myeloid (MHC-II) and functional markers (Ki67, IFNγ, PD1, ICOS and CD69) in CD14^+^CD4^+^ and CD14^-^CD4^+^ T cells in NSCLC as assessed by flow cytometry. Top panel show representative FACS image for each marker. Cells were gated on Live CD45^+^CD3^+^CD4^+^CD14^+/-^ cells. n=4 LUAD, n=3 LUSC and n=1 carcinoid. **\***\*, p<0.01; *, p<0.05, paired t-test between CD14^+^ and CD14^-^ T cell subsets. **C**. Kaplan-Meier survival curve showing overall survival of L-enriched patients in Sorin *et al.* with low (n=64) or high (n=47) CD14^+^CD4^+^ T cells infiltration in the tumor. *, p<0.05; Cox proportional hazards regression model (Wald test). **D.** Spatial neighborhood analysis of immune cells in Sorin *et al.* Left panel: pie charts of proportion of cell neighborhood (CN) clusters in patients with low (top) or high (bottom) CD14^+^CD4^+^ T cells infiltration in the tumor core. Right panel: volcano plots comparing the proportion of each CN clusters in the two compartments (stroma and tumor core) between patients with low (n=64) or high (n=47) CD14^+^CD4^+^ T cells infiltration in the tumor. A horizontal dashed line indicates -log10(0.05), Wilcoxon test. **E.** Barcode plot showing enrichment for TNF⍰ signaling pathway (198 genes) in tumors with high CD14^+^CD4^+^ T cells infiltration. The horizontal axis shows t-statistics for all genes comparing CD14^+^CD4^+^ high with low tumors, and the vertical bars show the ranks of the genes from TNF⍰ signaling set of the hallmark collection of MSigDB. The line shows enrichment of this pathway relative to random ordering. **F.** Schematic representative of coculture of HLADR^+^ cells labelled with membrane-APC dye with CD3^+^TCR⍰β^+^CD14^-^ T cells without or with TNF⍰ (top). Bar plot showing fold change in membrane-labelled CD14^+^CD4+ T cells after treatment with TNF⍰ compared with vehicle-treated cells (bottom). n=3.

We then examined the clinical relevance of CD14^+^CD4^+^ T cells infiltration in tumors. High infiltration of CD14^+^CD4^+^ T cells in L-enriched tumors (Sorin *et al*. n=111) was significantly associated with poorer overall survival (median survival 9 yrs vs 12.8 yrs in CD14^+^CD4^+^ high vs low tumors, respectively) (Fig 6C). Spatial neighborhoods analysis showed that tumors with high CD14^+^CD4^+^ T cell infiltration had a distinct cellular microenvironment, enriched in macrophage/T cell clusters (Fig 6D, Supp Fig 6C), consistent with the observation that CD14^+^CD4^+^ T cells reside in myeloid-rich niches (Fig 5D, E, F). Evaluation of the demographics and clinical characteristics of the patients in CD14^+^CD4^+^ low and high tumors show similar age, sex, BMI, smoking status and disease stages between the two groups. Histological patterns were slightly different with more acinar patterns in CD14^+^CD4^+^ low tumors and more solid patterns in CD14^+^CD4^+^ high tumors (Supp Table 3).

To evaluate whether the infiltration of CD14^+^CD4^+^ T cells could be associated with a specific tumor cell phenotype, we compared the transcriptional profiles of tumor cells in CD14^+^CD4^+^-high tumors and CD14^+^CD4^+^-low tumors. KEGG pathway enrichment analysis identified cytokine signaling pathways upregulated in tumors with high CD14^+^CD4^+^ T cells infiltration, indicative of an inflammatory microenvironment, while metabolic pathways were downregulated in tumors with high infiltration of CD14^+^CD4^+^ T cells (Supp Fig 6D). Specifically, the TNF⍰ signaling pathway was enriched in tumors with high infiltration of CD14^+^CD4^+^ T cells (Fig 6E). Given the known immunosuppressive effect of TNF⍰ on the tumor microenvironment^20^, we evaluated whether it could alter the properties of CD4^+^ T cells by enhancing T cell trogocytosis from myeloid cells. Trogocytosis assay showed that TNF⍰ increased the capture of membrane components by CD4^+^ T cells from HLA-DR^+^ cells by 1.7 to 1.9 fold (Fig 6F) compared to vehicle-treated cells. These results show that TNF⍰ present in the TME may enhance CD4^+^ T cell trogocytosis, enhancing the formation of myeloid-instructed T cells, a mechanism that may contribute to TNF⍰-mediated pro-tumorigenic properties.

## Discussion

Advances in cancer immunotherapy have greatly improved outcomes for lung cancer patients. However, not all patients respond to treatment^1,2^, indicating that alternative approaches to increase anti-cancer immunity are urgently needed. In-depth characterization of the composition and spatial organization of individual immune cells in the TME, and the signals that regulate them, is essential in identifying pathways to deplete pro-tumorigenic cells to bolster anti-tumor immunity. Here, we performed deep spatial proteomic and transcriptomic profiling of immune and tumor cells in NSCLC samples. We identified that neutrophil communities present at the border of the tumor front are associated with short-term patient survival in LUAD and LUSC. We show that myeloid-instructed CD14^+^ T cells are associated with short-term patient survival in LUAD. Tumors with high CD14^+^CD4^+^ T cell infiltration present a transcriptomic profile enriched in TNF⍰ signaling. We observed that TNF⍰ promotes CD4^+^ T cells trogocytosis from myeloid cells, augmenting the proportion of myeloid-instructed CD14^+^CD4^+^ T cells in co-culture assays. CD14^+^CD4^+^ T cells exhibit an atypical immune phenotype, suggesting that myeloid-instruction may alter their anti-tumor response.

*In situ* analysis of individual cells in solid tumors presents a clear advantage over single cell analysis of dissociated tissue. Notably, the architecture of the tissue is preserved, adding to the depth of knowledge on the role of cell communities in the anti-tumor immune response. In addition, this approach also avoids disproportionate recovery of different cell types, which is known to occur after tissue disruption^35^. Our comparison of the *in situ* TME in different histological subtypes of NSCLC revealed similarities and distinctions in the composition and organization of the TME of LUAD and LUSC. As described by others^11,24^, we observed a higher proportion of immune cells in the stroma and at the tumor border than in the tumor core. However, macrophages and CD8^+^ T cells were more predominant in the tumor core than the border and the stroma in both LUAD and LUSC. Myeloid cells were always observed in LUSC, whereas nearly half of the LUAD samples had no to little myeloid cell infiltration. Cell neighborhood analysis showed that immune cells were mostly present as clusters of single cell types or cells from similar lineages. Mixed myeloid and T-cell cellular neighborhoods were a distinctive feature of the LUSC TME, whereas B-cell clusters were only observed in LUAD and associated with better survival in LUAD, as described by Sorin *et al*^13^.

By profiling the TME in NSCLC, we were able to stratify LUAD and LUSC in different immune categories. M-enriched tumors had a distinct myeloid cell infiltration compared with LM-enriched tumors. While neutrophils were the predominant myeloid cell type in M-enriched tumors, macrophages constituted the bulk of the myeloid infiltrate in LM-enriched tumors. Neutrophils have a multi-faceted role in cancer^36,37^ but have often been associated with worse clinical outcomes in solid tumors, facilitating tumor cell extravasation and metastasis^38–40^. A high neutrophil-to**-**lymphocyte ratio has been associated with worse overall survival in many solid malignancies, including lung cancer^16,29^, consistent with our observation that tumors with clusters of neutrophils had worse survival.

Focusing on tumors with lymphoid cell infiltration, we observed that a subset of CD4^+^ and CD8^+^ T cells expressed the myeloid cell marker CD14, similar to the CD14^+^CD8^+^ and CD14^+^CD4^+^ T cells described in human liver^18^. Detailed analysis by Pallett *et al.* demonstrated that CD14^+^ T cells were myeloid-instructed T cells, in which CD8^+^ T cells acquired cellular myeloid materials from CD14^+^ myeloid cells in a contact-dependent manner. Exposure to bacterial LPS promoted the transfer of myeloid materials to T cells, with the presence of bacteria reshaping the local immunity and inducing the expansion of CD14^+^CD8^+^ T cells^18^. The lung is host to a diverse range of commensal bacteria^41^ and increased bacterial colonization is observed in lung tumors^42–45^. Further studies would be required to investigate the relationship between CD14^+^ T cells, the tumor microbiota and the implication for response to immunotherapy.

Importantly, we observed that patients with tumors with a high infiltration of CD14^+^ expressing CD4^+^ T cells had poor survival, pointing to CD14^+^CD4^+^ T cells being unable to support anti-tumor immunity. T cell trogocytosis is a known phenomenon that has been shown to involve antigen-presenting cells^46^ or B cells^47^ leading to the transfer of a number of immune regulatory molecules such as HLA-G, PDL1, OX40L that can reduce anti-tumor immunity. We demonstrated that CD14^+^CD4^+^ T cells can form through trogocytosis and that in NSCLC, these myeloid-instructed CD14^+^CD4^+^ T cells were more frequent than CD14^+^CD8^+^ T cells. CD14^+^CD4^+^ T cells had an unusual phenotype expressing markers of T cell activation (IFNγ, Ki67, CD39) but low levels of PD1, ICOS and CD69. Trogocytosis has been described as an antigen-specific, TCR affinity-dependent process^48,49^. Our observation of a clonal expansion of CD14^+^CD4^+^ T cells compared to CD14^-^CD4^+^ T cells, suggests that antigen-specific T cells in tumors may be more prone to trogocytosis. We found that TNF⍰ increased CD4^+^ T cell trogocytosis from HLA-DR^+^ myeloid cells. TNF⍰ can exert both tumor-promoting and tumor-suppressing activities^20^, but combination drug studies show that inhibition of TNF⍰ in combination with immune checkpoint inhibitors increases treatment response^50^, suggesting a predominant role for TNF⍰ in promoting tumour growth. TNF⍰ exerts its pro-tumorigenic activity in multiple ways, including the recruitment of myeloid and myeloid-derived suppressor cells to the TME and maintaining regulatory T cell function^20,51^. Here we identify a possible new mechanism by which TNF⍰ exerts its pro-tumor activity, by facilitating T cell trogocytosis, thereby altering their anti-tumor properties. Future work would be needed to evaluate whether CD14^+^CD4^+^ T cells are depleted in tumors treated with TNF⍰ inhibitors, such as Infliximab, and determine the relative contribution of this process to the overall pro-tumorigenic activities of TNF⍰.

Overall, our study demonstrates the importance of undertaking a comprehensive analysis of the tumor ecosystem with hi-plex *in situ* molecular approaches to uncover novel cell types and decipher spatial relationships between immune and tumor cells. Our study revealed myeloid-instructed T cells in the TME may participate in tumor immune escape. Targeting the recruitment or formation of myeloid-instructed T cells to the tumor core, such as with the TNF⍰ inhibitor Infliximab, may provide novel therapeutic avenues for lung cancer patients.

## Methods

### Human NSCLC cohort

Written informed consent was obtained from all patients by the Victorian Cancer Biobank prior to inclusion in the study, according to protocols approved by the WEHI Human Research Ethics Committee (HREC, 10/04LR). Human tonsil FFPE tissues were obtained were used for antibody validations. NSCLC specimens and associated clinical data were obtained from the Victorian Cancer Biobank from treatment-naïve, surgically resected early-stage lung cancer patients (Stage I to IIIa). Patients were classified as ever-smokers (current and ex-smokers who had quit smoking at least 1 year prior to sample collection) or never-smokers (lifetime smoking of less than 100 cigarettes). Only patients with an overall survival (OS) of less than 3 years post-surgery and with OS of more than 6 years post-surgery were included in the study. Human tissues from non-cancer patients were obtained from deceased organ donors at the time of organ acquisition for transplantation through the Australian Donation and Transplant Biobank^52^ (HREC/4814/Austin-2019). All donors were free of cancer, hepatitis B, and hepatitis C, and were HIV negative. PBMC used for the trogocytosis experiments were collected from healthy donors through the Victorian Blood Donor Registry (HREC 10/04LR). Patient and organ donor details are described and summarized in Supplementary Table 1 and 2. Three formalin-fixed paraffin-embedded (FFPE) serial sections (4µm thickness) were cut for H&E staining, MIBI proteomic experiment on gold-coated slides (Ionpath #567001), and for GeoMX transcriptomic experiment.

### Validation of 38-plex antibody panel for MIBI

Rare earth polymer conjugated antibodies were purchased from Ionpath (Supp Table 3) or conjugated to metal polymers according to manufacturer protocol (Ionpath). Briefly, 100µg of carrier-free antibody was reduced with 4mM TCEP. Partially reduced antibody was then incubated at 37°C for 1hr with the metal-loaded polymer. The antibody concentration was quantified by measuring the IgG absorbance at 280nm on the NanoDrop. The conjugated antibody was eluted to obtain a final concentration of 0.5mg/ml. All antibodies were first validated by immunohistochemistry on NSCLC tissue sections. Panels of conjugated antibodies were then evaluated on human tonsils to ensure specificity and the absence of isobaric and isotopic overlap. Antibodies were then titrated on lung cancer tissues before being pooled in a 38-plex panel for staining of the complete cohort (Supp Table 4 and Supp Fig 1A).

### Slide staining and scanning on MIBI

Slides were processed as per manufacturer protocol and stained with our bespoke 38-plex antibody panel. Antigen retrieval was performed with Tris EDTA pH9 (Agilent, #S236784-2) using the PT Link Module (Thermo Scientific), sections blocked with 10% donkey serum and stained overnight with 36 conjugated antibodies. Slides were then washed in TBS-0.1% Tween and stained for 30 min with 2 antibodies (Supp Table 4) before being dehydrated and post-fixed with 2% glutaraldehyde. Slides were stored in a vacuum chamber before scanning. Fields of view of 400×400µm (1024×1024 pixels) were acquired at a dwell time of 1ms. The mass-spec pixel data generated is automatically converted into TIFF images. MIBI images were pre-processed for isobaric correction using MIBI/O (Ionapth) and visualized on QuPath^22^. Adjacent FOVs were stitched together using a groovy script plugin in QuPath^53^.

### Single-cell segmentation

Cell segmentation was performed using a QuPath extension applying Mesmer^23,54^. The dsDNA channel was used as the nuclei features and the matrix addition of the median normalized MHC I channel and the panCK channel was used as the membrane features. Segmented cells with a nuclei area of <50 pixels and cells touching the border of images were removed.

### Single-cell phenotyping

Cell classification was performed using the supervised method XGBoost. We manually annotated 52,719 cells using in-house developed tools on QuPath. Cells were annotated as epithelial cells, stromal cells, B cells, CD4^+^ T cells, CD8^+^ T cells, Treg cells, NK cells, *γδ*T cells, macrophages, dendritic cells, neutrophils, mast cells, or others. Annotations were assigned to cells based on the shape and intensity of the markers within the cell boundaries visible on the images. All markers were used to train the model. Each marker was measured in four compartments: nucleus, cytoplasm, membrane, and the entire cell. In each cell compartment, the measurements calculated were mean, standard deviation, and the 0.7, 0.8, 0.9, 0.95, 0.96, 0.97, 0.98, 0.99, 0.995, 0.999 quantiles. A polynomial feature generation of degree two was used between markers expected to be co-expressed. The combinations were generated from pools of markers expected to be co-expressed. These features were generated on cell mean measurements of the markers. In total 1713 features were used to train the model. To compensate for the imbalanced data, we used the synthetic minority oversampling technique (SMOTE). To select the hyper-parameters, we used the Bayesian search cross validation (https://scikit-optimize.github.io/stable/modules/generated/skopt.BayesSearchCV.html). In assessing the results of the training confusion matrices, scores such as f1, balanced accuracy, ROC AUC score, were used.

### Tissue compartment classification immune subgroups

Twelve different pixel classifiers were trained in QuPath and applied to our cohort of 64 patients to delimit the tumor and stroma area. For spatial analysis, three compartments were created: tumor core, border and stroma where the border was defined as 40µm inside and outside the tumor area. Patients were grouped into myeloid-, lymphoid-myeloid- or lymphoid-enriched (M-, LM-, L-enriched) after an unsupervised k mean (k=3) clustering of the proportions of lymphoid and myeloid cells in each sample.

### Cell neighborhood clustering of immune cells

Cell neighborhood (CN) clusters of immune cells were defined by first determining the 10 closest neighbors (excluding itself) of every cell. The cell type frequencies were determined and converted into proportions. These proportions were used in a K means clustering with k=7. Cell type proportions have been z-score normalized within each cluster.

### Statistical analysis

Statistical analyses were performed using R (2023.06.1+524). Boxplots present median +/- standard deviation. Non-normal data were compared between two groups using Wilcoxon rank sum test. Samples with less than 5 cells of the cell type of interest were excluded from the analysis. The significance between two groups in Kaplan Meier survival curves was calculated using log-rank (Mantel-Cox) test. In the heatmaps, percentages are calculated for each patient and then z-score normalized across all patients using the scale function with default parameters in R.

### Sorin *et al.* and Enfield *et al.* analysis

Data from Sorin *et al.*^13^ were available on Zenodo (https://doi.org/10.5281/zenodo.7383627). Data from Enfield *et al.*^29^ were kindly provided by the authors under a Data Transfer Agreement. The same k-means clustering (k=3) used in our cohort was applied to both datasets using their cell phenotyping grouping of myeloid and lymphoid cells in each patient. To delimitate the ‘in’ and ‘out’ tumor area in Sorin et al., a cell neighborhood analysis was performed using **imcRtools** package^55^. Interaction graphs based on the cell’s locations were constructed using distance-based expansion with radius 30, and then neighbors were aggregated by cell type (top hierarchical level: Cancer, Endothelial cells, Immune cells and Others) and k-means clustering was performed with k=4. All the cells within two clusters mainly composed of tumor cells were classified as inside the tumor area.

#### Defining CD14^+^ cells

To define cells as CD14 positive we first expanded nuclear boundaries provided by Sorin et al. by 3 pixels using functionality of the **EBImage** R package^56^ and re-calculated the mean intensity of CD14 for each cell. To define CD14-positive cells among non-monocytes, we employed a data-driven classification approach based on CD14 expression profiles in annotated monocyte populations. Classical monocytes (CD14□CD16□) were used as the positive reference group, and non-classical monocytes (CD14□CD16□) as the negative reference group, reflecting high and low CD14 expression levels, respectively.

For each group, we estimated the empirical density of CD14 expression using kernel density estimation (KDE) with gaussian kernel. The log-density ratio (LDR) was then computed at each expression value for all patients together. The LDR reflects the relative likelihood that a given CD14 expression value originates from an expected CD14 positive cell (classical monocyte) rather than a non-classical monocyte, which we expect to be CD14 negative. To account for the imbalance in group sizes (with classical monocytes substantially outnumbering non-classical monocytes), we adjusted the classification threshold using the log of the observed group size ratio. This choice reflects a Bayesian decision-theoretic framework with prior probabilities proportional to group sizes and a 0–1 loss function, yielding a threshold that minimizes expected misclassification under class imbalance^57^. Cells with CD14 expression levels yielding an LDR greater than this threshold were classified as CD14 positive.

#### Calculating distances to CD14^+/-^CD4^+^ T cells

We used the *minDistToCells* function of the **imcRtools** package to calculate minimal distance to cells of interest (CD14+/- CD4+ T cell). Based on this distance, cells that are further than 50 units away from the cells of interest were excluded from the neighborhood analysis presented in Fig. 5 E and F. Significant differences in immune cell type composition of neighborhoods between CD14^+^ and CD14^-^ CD4+ T cell were assessed using a *propeller* test from the **speckle** Bioconductor R package (version 1.2)^58^.

#### Cell-cell interactions

Cell-cell interactions were counted and tested using the *testInteractions* function of the **imcRtools** package. Cancer cells were excluded prior to building the cell-interaction graph with distance-based expansion with radius 30μm.

#### Cell neighborhood clustering of immune cells

Cell neighborhood clustering of immune cells in Sorin *et al.* was performed as per our cohort by using k-means clustering with k=6.

### Human lung and tumor tissue dissociation

Lung and tumor samples were either processed immediately or held intact for a maximum of 16□h at 4°C in DMEM/F12 media (Gibco) supplemented with 1□mg/mL of penicillin and streptomycin (Invitrogen). Surgical samples were minced then digested for 1□h at 37°C with 2□mg/mL collagenase I (Worthington, #LS004197) and 200□U/mL deoxyribonuclease (Worthington, #LS002140) in 0.2% D-glucose (Sigma) in DPBS (Gibco), as previously described ^5^. The cell suspension was filtered through 100□µm strainer (Falcon) and washed with 2% FCS-PBS, followed by red blood cell lysis and further washing with 2% FCS-PBS to obtain a single cell suspension. Single cell suspensions were either stained immediately for flow cytometry analysis or cryopreserved in 90% FCS:10% DMSO and stored in liquid nitrogen prior to staining.

### Flow cytometry

Cells were stained with fixable viability dye Zombie NIR (Biolegend) for 15□min. Cells were washed with 2%FCS-PBS, then blocked with anti-CD16/32 FCRγ antibody (WEHI antibody facility) for 10□min. Cells were stained with extracellullar antibodies for 30□min at 4°C (Supp Table 4). Cells were then fixed and permeabilized with Foxp3/Transcription Factor Staining Kit (eBiosciences) and stained with intracellular antibodies for 30□min at 4°C (Supp Table 5). Samples were resuspended in FACS buffer (2% FCS in PBS) and data acquired using an Aurora spectral unmixing cytometer (CyTEK). Interrogation of singlet and doublet cells was performed by staining cells with extracellular antibodies for 30 mins 4°C (Supp Table 6) then incubated with Hoechst 33342 (5µg/ml) for 30min at °C. Samples were acquired using a BD Symphony flow cytometer. Data were analyzed using FlowJo (Tree Star).

### Image Cytometry

Real time images were captured using BD FACSDiscover™ S8 Cell Sorter running BD FACSChorus™ acquisition software (BD Biosciences, San Jose). The instrument contains 5 lasers with 6 spatially separated laser intercepts (the blue laser is split to support full-spectrum detection at one intercept and BD CellView™ imaging detection at a second intercept. Live cells were gated on CD45-PE^+^CD3-BV570^+^CD4-RB545^+^. The images for each cell were captured in real-time with the following detectors and bandpass filters, light-loss, forward scatter, side scatter (488/15), Image Channel 1 (534/46, CD4), and image channel 3 (788/225, CD14). Co-localization of CD4 and CD14 were determined using the imaging data and the following image analysis, Max Intensity (CD14 BB700) vs Correlation (CD4 RB545/CD14 BB700) plot.

### Trogocytosis

The protocol was adapted from Daubeuf *et al*.^59^. In brief, peripheral blood mononuclear cells (PBMCs) were isolated from healthy donors and stained with an antibody cocktail (Supp Table 7) to sort CD3□TCR⍰β^+^CD14□ and CD3□TCR⍰β^-^CD14□HLA-DR□ cell populations. Following cell sorting, each subset was independently labelled with 1 μg/mL of biotinylated membrane dye (EZ-Link™ Sulfo-NHS-LC-Biotin, Catalog #21335) in PBS for 10 minutes at 25°C. An equal volume of fetal calf serum (FCS) was then added, and the cells were incubated for an additional 10 minutes at 4°C. Subsequently, the biotin-labeled cells were co-cultured with unlabelled cells at a 1:1 ratio (total 1 × 10□ cells) at 37°C for 4 hours. The cells were acquired using NovoCyte Penteon Flow cytometer and analyzed using FlowJo 10.9.0. For TNF-⍰ treatment, 100ng/µL stock solution of recombinant human TNF⍰ (catalogue # ab9642) was prepared in dPBS. Appropriate volume of the stock solution was added to 4 hrs co-culture experiment described above so that the final concentration of TNF⍰ was 10 or 80ng/mL.

### TCR sequencing

CD45^+^CD3^+^CD4^+^ CD14^+^or CD14^-^ T cells were sorted on the Aria Fusion (BD) (Supp Table 8) and DNA extracted using DNeasy Blood & Tissue kit (Qiagen). DNA Integrity was measured on the Tapestation 4200 (Agilent). Samples with a DNA Integrity Number > 8.5 were sent for TCRβ sequencing by ImmunoSeq (Adaptive Biotechnologies, Seattle). TCR reads and sequences were analyzed on the Immunoseq data analysis platform (Adaptive Biotechnologies).

### Oncogenic driver mutation sequencing

*EGFR, KRAS* and *STK11* mutation analyses were performed on fresh frozen tumor samples. Genomic DNA was prepared using the Qiagen DNeasy Blood and Tissue kit (Qiagen #69504) according to manufacturer’s instructions. Sequencing was performed using a 2-step PCR approach for targeted sequencing. Unique primer sequences for *EGFR, KRAS* and *STK11* that contain a common overhang sequence at the 5’ end were used to amplify *EGFR, KRAS* and *STK11* exonic regions of interest. Cycling conditions were initial denaturation at 96°C for 5min, 18 cycles of denaturation at 96°C for 30sec, annealing at 57°C for 30sec, and elongation at 72°C for 30sec, followed by a final elongation step at 72°C for 5min. PCR amplicons were purified using 1.0x Ampure Beads (Beckman Coulter) prior to the second PCR to introduce adaptor sequences and 8 bp dual-index barcodes for multiplexing. Cycling conditions were initial denaturation at 95°C for 3min, 24 cycles of denaturation at 95°C for 15sec, annealing at 60°C for 30sec, and elongation at 72°C for 30sec, followed by a final elongation step at 72°C for 7min. The barcoded PCR product size was assessed by agarose gel electrophoresis, purified using 0.8x Ampure Beads and quantified by Qubit fluorometry (ThermoFisher Scientific). PCR products from up to 96 samples were pooled equimolar and sequenced on a MiSeq (Illumina). Read pairs were demultiplexed according to the index sequences using bcl2fastq, and subsequently processed by Trim_galore (https://www.bioinformatics.babraham.ac.uk/projects/trim_galore/) to remove adaptors and low-quality reads. The clean reads were aligned using the https://sarah-d.shinyapps.io/crispr-indel/, to WT sequence from NCBI RefSeq Genome database GRCh38. Primer sequences are detailed in Supplementary Table 9. For *STK11,* we screened mutations in *STK11* exon 1 and 2, as these are associated with worse survival in NSCLC patients^60^. We investigated the presence of previously described alterations in exon 1 *Q37Ter* and *K44Ter* and screened for mutations in all of exon 2. For *EGFR* we evaluated *ex19-del*, *L858R*, and *T790M* mutations. For *KRAS*, we assessed mutations in G12 and G13 codons.

### GeoMx data collection

FFPE sections from 64 NSCLC samples were sent to the NanoString Centre of Excellence in Spatial Biology (Griffith University) for sample processing and data collection on the GeoMx platform (Nanostring). Slides were baked at 60°C for 90 min. and deparaffinised at 72°C for 30min, Heat-induced epitope retrieval for 20 min and Proteinase K treatment (1µg/mL) for 15 min using the automated LEICA BOND Rx system. Immunofluorescent staining was performed using the morphology antibodies from Nanostring Solid Tumor Morphology kit: SYTO13 (nucleus, 1:10 dilution), PanCK-AF532 (tumor cells, 1:40 dilution) and CD45-AF594 (immune cells, 1:40 dilution). For in-situ hybridization for next generation sequencing, tissue sections were then hybridized with Whole Transcriptome Atlas (WTA; Lot# HWTA21003) probes for 16 hours at 37°C in a humidity-controlled chamber. Two segments were collected: PanCK positive cells and CD45 positive cells, with a minimum of 200 cells collected for each individual area of illumination (AOI). A total of 345 AOIs were collected.

### GeoMx data analysis

NanoString GeoMx DSP data was processed with GeoMx DSP Data Analysis Suite (https://nanostring.com/products/geomx-digital-spatial-profiler/geoscript-hub/). All downstream analysis was performed in the R statistical programming language (version 4.3.2) using predominantly Bioconductor packages (version 3.18). Further quality control was performed using the **standR** Bioconductor R package version 1.6^61^. Library sizes were normalized with the TMM method^62^, as implemented in the *geomxNorm* function of the **standR** package. Count data were transformed to log2-counts per million, voom precision weights were estimated and linear models for differential expression analysis were fitted using **edgeR**’s *voomLmFit* function with additional cyclicloess normalization (**edgeR** version 4.0)^63,64^. For the comparison of LUAD and LUSC cancer types, age and sex were included as covariates in the linear model. Differential expression was assessed using moderated t-statistics^65^ with robust empirical Bayes variance estimation^66^. Genes with a false discovery rate < 0.05 and log-fold-change threshold > 1.1 were defined as significantly differentially expressed using TREAT^67,68^. For LUAD M vs LLM comparison, the cancer sub-types, age and smoking status were included in the linear model. Genes were ranked using p-values obtained from moderated t-statistics. The top ranked genes with a log-fold change < -0.5 and > 0.5 are highlighted in the mean-difference plot. For the comparison of CD14^+^-high vs CD14^+^-low tumors, CD14 status was included in the linear model together with smoking status. CD14 status was determined as high/low using the MIBI data and defined as areas of illumination (AOI) with greater/smaller percentage of CD14 positive cells than the median value across all AOI. Gene set testing to test for enrichment of REACTOME gene sets, Gene Ontology terms, KEGG pathways and Hallmark gene sets from the Molecular Signatures Database (MSigDB)^69^ was performed using the *camera* function in **limma** Bioconductor R package (version 3.58)^70,71^. For Fig 5G, Fig 3F, Supp Fig 4F and Supp Fig 6F, genes from selected pathways were highlighted using barcode plots, where the x-axis represents ordered moderated t-statistics from the testing procedure described above. Significant differences in proportions of cells with higher expression of functional markers between CD14^+^ and CD14^-^ were assessed using a *propeller* test from the **speckle** Bioconductor R package (version 1.2)^58^.

## Data Availability

The sequence data generated in this study are publicly available in Gene Expression Omnibus (GEO) at GSEXXXX (TBC) and within the article and its supplementary data files. Codes are available at https://github.com/WEHI-labatlab/

## Acknowledgements

We thank WEHI facilities including Histology (Emma Pan and Ellen Tsui) and Flow cytometry (Simon Monard) for expert advice and performing experiments. We are grateful to the Victorian Cancer Biobank and all lung cancer patients who participated in this study. Real time images of the cells were captured using a BD FACSDiscover™ S8 Cell Sorter located at BD Melbourne, Australia with the assistance of the BD Biosciences ANZ applications team. The MIBI studies were supported by the Australian Cancer Research Foundation Program for Resolving Cancer Complexity and Therapeutic Resistance. C.M. was supported by a bequest from the late Margaret Anne Ireland and funds from the Harry Secomb Trust-Perpetual Impact. B.P. is supported by an NHMRC Investigator Grant (GNT1175653). The MIBI studies were supported by the Australian Cancer Research Foundation Program for Resolving Cancer Complexity and Therapeutic Resistance. This work is supported by a NHMRC Ideas Grant (GNT1182155), a Lyn Williams research grant, Jenny Tatchell research grant and by funds from the Operational Infrastructure Support Program provided by the Victorian Government and NHMRC IRIISS (Independent Research Institutes Infrastructure Support Scheme) Grant.

## Supplementary Figure legends

***Supp Fig. 1. Validation of 38-plex antibody panel used in MIBI analysis of NSCLC samples.* A.** Representative images of MIBI staining for each indicated marker. **B.** UMAP of all cell phenotypes generated using the mean marker measurement of each phenotypic marker in all cells in all samples. The colors reflect the phenotype assigned by the XGBoost model.

***Supp Fig 2. Characterization of immune and tumor cells in resected LUAD and LUSC samples.* A.** Mean-difference plot highlighting differentially expressed genes in tumor cells (AOI panCK^+^ cells) between LUAD and LUSC samples. **B.** Kaplan-Meier survival curve showing relapse-free survival in LUAD patients based on the level of expression of different markers in tumor cells, as measured by analysis of MIBI data or pathologist scoring for PDL1. Log-rank (Mantel-Cox) test not significant for all comparison. Except for PDL1, the cut-off used to separate low and high expression is the median. **C** and **D.** Boxplots showing the localization of all immune cells (**C**) and indicated immune cell types (**D**) in the different tissue compartments (tumor core, border and stroma) in LUAD and LUSC patients. ****, p<0.0001; ***, p<0.001; **, p<0.01; *, p<0.05 comparing the different compartments, Wilcoxon test. n= 44 LUAD, n=20 LUSC. **E.** Volcano plot comparing the proportion of each immune cell subtypes in the whole tissue in LUAD and LUSC samples. A horizontal dashed line indicates -log10(0.05), Wilcoxon test.

***Supp Fig 3: Characterization of immune-enriched tumor subgroups in LUAD and LUSC.* A.** Heatmaps showing the k-mean (k=3) clustering stratification of the TME in LUSC samples in our cohort (i) and in Enfield *et al.* cohort (ii). Both LUSC cohorts are stratified in 2 immune subgroups lymphoid/myeloid-enriched (LM-enriched; n=15 (i), n=10 (ii)) or myeloid-enriched (M-enriched; n=5 (i), n=13 (ii)). **B.** Kaplan-Meier survival curve showing relapse-free survival (RFS) of LLM-enriched LUAD patients with low or high B cell infiltration in the tumor core. n=12 low B cell-infiltrated tumors; n=6 high B cell-infiltrated tumors. *, p<0.05; log-rank (Mantel-Cox) test. The cut-off used to separate low and high expression is the lower and upper quantile. **C.** Box plot showing the proportion of myeloid cells in each tissue compartment in LUAD and LUSC tumors in the M-enriched immune subgroups. Data show median ± SD. *, p<0.05 comparing the different compartments, Wilcoxon test. n=9 LUAD, n=5 LUSC **D.** Heatmap showing the mean expression of each marker z-score normalized across all the cell types in the three spatial compartments in M-enriched LUAD tumors. *, p<0.05, Wilcoxon test. n=9 M-enriched LUAD. **E.** Volcano plots comparing the proportion of each CN cluster in the three compartments between M and LM-enriched LUSC samples. A horizontal dashed line indicates -log10(0.05), Wilcoxon test. **F.** Kaplan-Meier survival curve showing overall survival in the three immune subgroups of LUAD patients n=9 M-enriched, n=16 LM-enriched, n=19 L-enriched. *, p<0.05 comparing LM-enriched with M-enriched tumors; Cox proportional hazards regression model (Wald test). **G.** Representative image showing the infiltration of mast cells in the M-enriched LUAD tumor (MH1260) with long relapse-free survival. Scale bar: 50µm. **H.** Kaplan-Meier survival curve showing overall survival in the two immune subgroups of LUSC patients n=5 M-enriched, n=15 LM-enriched. *, p<0.05 comparing LM-enriched with M-enriched tumors; Cox proportional hazards regression model (Wald test).

***Supp Fig 4: Characterization of CD14^+^ T cells in LUAD and LUSC tumors.* A.** Representative image showing CD4^+^, CD8^+^ T cells and macrophages positive and negative for CD14 in LUAD tumor (AH0353). Scale bar: 50µm. **B.** Proportion of immune cell types expressing CD14 in LUSC as determined by MIBI. Data show median ± SD. n=20 LUSC. **C.** Proportion of immune cell types expressing CD14 in LUAD Sorin *et al.*. Data show median ± SD. n=416 LUAD. **D.** Box plot showing the proportion of lymphoid cell subsets expressing CD14 in LUSC as determined by MIBI. Data show median ± SD. n=20 LUSC. **E.** Boxplot showing the localization of CD14^+^CD4^+^ T cells in the different tissue compartments (tumor core, border and stroma) in LUAD and LUSC patients. **, p<0.01; *, p<0.05, Wilcoxon test. n=41 LUAD, n=20 LUSC. **F.** Representative flow cytometric gating strategy used to identify live CD45^+^CD3^+^CD14^+^CD4^+^ T cells in non-small cell lung cancer (MH0069). **G.** Representative flow cytometry histograms showing the percentage of single or doublet CD14^+^CD4^+^ T cells after Hoestch staining (MH0061).

***Supp Fig 5: Characterization of myeloid-instructed CD14^+^CD4^+^ T cells and their microenvironment.* A.** Frequency of paired productive TCR sequences (top 20) in paired CD14^+^CD4^+^ and CD14^-^CD4^+^ T cells. n=2 LUAD and n=2 LUSC. The right panels show frequency of the top 20 productive TCR sequences in CD14^+^CD4^+^ and CD14^-^CD4^+^ T cells from individual LUAD (MH0059 and MH0069) and LUSC samples (MH0061 and MH0006). **B.** Correlation plots between the proportion of CD14^+^CD4^+^ T cells and classical macrophages (Cl Mac), alternative macrophages (Alt Mac), non-classical monocytes (non-Cl Mo), CD14^-^CD4^+^ T cells, CD14^-^CD8^+^ T cells, CD14^+^CD8^+^ T cells, NK cells, B cells, neutrophils, mast cells and dendritic cells (DC) in Sorin *et al.* L-enriched subgroup. Spearman rank correlation. n=113 LUAD.

***Supp Fig 6: Characterization of tumors with high infiltration of CD14^+^CD4^+^ T cells.* A.** Mean fluorescence intensity (MFI) expression of LAG3, OX40 and CTLA4 in CD14^+^CD4^+^ and CD14^-^CD4^+^ T cells in NSCLC as assessed by flow cytometry. Top panel shows a representative flow histogram for each marker. Cells were gated on Live CD45^+^CD3^+^CD4^+^CD14^+/-^ cells. n=4 LUAD, n=3 LUSC and n=1 carcinoid. **\***\*, p<0.01; *, p<0.05, paired t-test between CD14^+^ and CD14^-^ T cell subsets. **B.** Left panel: expression of CD103 in CD14^+^CD4^+^ and CD14^-^CD4^+^ T cells as determined by MIBI. *, p<0.05, paired t-test. Data show median ± SD. n=42 LUAD. Right panel: Mean fluorescence intensity (MFI) expression of CD103 in CD14^+^CD4^+^ and CD14^-^CD4^+^ T cells in NSCLC as assessed by flow cytometry. The top panel shows a representative flow histogram. n=4 LUAD, n=3 LUSC and n=1 carcinoid. Paired t-test between CD14^+^ and CD14^-^ T cell subsets. **C.** Heatmap showing six immune cell neighborhoods (CN) detected in Sorin *et al.* samples, and the cell type composition for each cluster. **D.** KEGG pathway analysis showing selected pathways with FDR< 0.05. n=6 low CD14^+^CD4^+^-infiltrated tumors (14 FOVs); n=3 high CD14^+^CD4^+^-infiltrated LLM-enriched tumors (8 FOVs).

## References

1. Langer, C.J., Gadgeel, S.M., Borghaei, H., Papadimitrakopoulou, V.A., Patnaik, A., Powell, S.F., Gentzler, R.D., Martins, R.G., Stevenson, J.P., Jalal, S.I., et al. (2016). Carboplatin and pemetrexed with or without pembrolizumab for advanced, non-squamous non-small-cell lung cancer: a randomised, phase 2 cohort of the open-label KEYNOTE-021 study. Lancet Oncol. 17, 1497–1508. 10.1016/s1470-2045(16)30498-3.

2. Gandhi, L., Rodríguez-Abreu, D., Gadgeel, S., Esteban, E., Felip, E., Angelis, F.D., Domine, M., Clingan, P., Hochmair, M.J., Powell, S.F., et al. (2018). Pembrolizumab plus Chemotherapy in Metastatic Non–Small-Cell Lung Cancer. N. Engl. J. Med. 378, 2078– 2092. 10.1056/nejmoa1801005.

3. Rizvi, H., Sanchez-Vega, F., La, K., Chatila, W., Jonsson, P., Halpenny, D., Plodkowski, A., Long, N., Sauter, J.L., Rekhtman, N., et al. (2018). Molecular Determinants of Response to Anti–Programmed Cell Death (PD)-1 and Anti–Programmed Death-Ligand 1 (PD-L1) Blockade in Patients With Non–Small-Cell Lung Cancer Profiled With Targeted Next-Generation Sequencing. JCO 36, 633–641. 10.1200/jco.2017.75.3384.

4. Hellmann, M.D., Nathanson, T., Rizvi, H., Creelan, B.C., Sanchez-Vega, F., Ahuja, A., Ni, A., Novik, J.B., Mangarin, L.M.B., Abu-Akeel, M., et al. (2018). Genomic Features of Response to Combination Immunotherapy in Patients with Advanced Non-Small-Cell Lung Cancer. Cancer Cell 33, 843–852.e4. 10.1016/j.ccell.2018.03.018.

5. Weeden, C.E., Gayevskiy, V., Marceaux, C., Batey, D., Tan, T., Yokote, K., Ribera, N.T., Clatch, A., Christo, S., Teh, C.E., et al. (2023). Early immune pressure initiated by tissue-resident memory T cells sculpts tumor evolution in non-small cell lung cancer. Cancer Cell. 10.1016/j.ccell.2023.03.019.

6. Angelova, M., Mlecnik, B., Vasaturo, A., Bindea, G., Fredriksen, T., Lafontaine, L., Buttard, B., Morgand, E., Bruni, D., Jouret-Mourin, A., et al. (2018). Evolution of Metastases in Space and Time under Immune Selection. Cell 175, 751–765.e16. 10.1016/j.cell.2018.09.018.

7. Kirchhammer, N., Trefny, M.P., Maur, P.A. der, Läubli, H., and Zippelius, A. (2022). Combination cancer immunotherapies: Emerging treatment strategies adapted to the tumor microenvironment. Sci Transl Med 14, eabo3605. 10.1126/scitranslmed.abo3605.

8. Bruni, D., Angell, H.K., and Galon, J. (2020). The immune contexture and Immunoscore in cancer prognosis and therapeutic efficacy. Nat. Rev. Cancer 20, 662–680. 10.1038/s41568-020-0285-7.

9. Lambrechts, D., Wauters, E., Boeckx, B., Aibar, S., Nittner, D., Burton, O., Bassez, A., Decaluwé, H., Pircher, A., Eynde, K., et al. (2018). Phenotype molding of stromal cells in the lung tumor microenvironment. Nat Med, 1–19. 10.1038/s41591-018-0096-5.

10. Guo, X., Zhang, Y., Zheng, L., Zheng, C., Song, J., Zhang, Q., Kang, B., Liu, Z., Jin, L., Xing, R., et al. (2018). Global characterization of T cells in non-small-cell lung cancer by single-cell sequencing. Nat Med, 1–17. 10.1038/s41591-018-0045-3.

11. Lavin, Y., Kobayashi, S., Leader, A., Amir, E.D., Elefant, N., Bigenwald, C., Remark, R., Sweeney, R., Becker, C.D., Levine, J.H., et al. (2017). Innate Immune Landscape in Early Lung Adenocarcinoma by Paired Single-Cell Analyses. CELL 169, 750–757.e15. 10.1016/j.cell.2017.04.014.

12. Leader, A.M., Grout, J.A., Maier, B.B., Nabet, B.Y., Park, M.D., Tabachnikova, A., Chang, C., Walker, L., Lansky, A., Berichel, J.L., et al. (2021). Single-cell analysis of human non-small cell lung cancer lesions refines tumor classification and patient stratification. Cancer Cell, 1–29. 10.1016/j.ccell.2021.10.009.

13. Sorin, M., Rezanejad, M., Karimi, E., Fiset, B., Desharnais, L., Perus, L.J.M., Milette, S., Yu, M.W., Maritan, S.M., Doré, S., et al. (2023). Single-cell spatial landscapes of the lung tumour immune microenvironment. Nature, 1–7. 10.1038/s41586-022-05672-3.

14. Desharnais, L., Sorin, M., Rezanejad, M., Liu, B., Karimi, E., Atallah, A., Swaby, A.M., Yu, M.W., Doré, S., Hartner, S., et al. (2025). Spatially mapping the tumour immune microenvironments of non-small cell lung cancer. Nat. Commun. 16, 1345. 10.1038/s41467-025-56546-x.

15. Casanova-Acebes, M. x000ED a, Dalla, E., Leader, A.M., LeBerichel, J., Nikolic, J., Morales, B.M., Brown, M., Chang, C., Troncoso, L., Chen, S.T., et al. (2021). Tissue-resident macrophages provide a pro-tumorigenic niche to early NSCLC cells. Nature, 1–27. 10.1038/s41586-021-03651-8.

16. Templeton, A.J., McNamara, M.G., Šeruga, B., Vera-Badillo, F.E., Aneja, P., Ocaña, A., Leibowitz-Amit, R., Sonpavde, G., Knox, J.J., Tran, B., et al. (2014). Prognostic Role of Neutrophil-to-Lymphocyte Ratio in Solid Tumors: A Systematic Review and Meta-Analysis. JNCI: J. Natl. Cancer Inst. 106, dju124. 10.1093/jnci/dju124.

17. Barry, S.T., Gabrilovich, D.I., Sansom, O.J., Campbell, A.D., and Morton, J.P. (2023). Therapeutic targeting of tumour myeloid cells. Nat. Rev. Cancer 23, 216–237. 10.1038/s41568-022-00546-2.

18. Pallett, L.J., Swadling, L., Diniz, M., Maini, A.A., Schwabenland, M., Gasull, A.D., Davies, J., Kucykowicz, S., Skelton, J.K., Thomas, N., et al. (2023). Tissue CD14+CD8+ T cells reprogrammed by myeloid cells and modulated by LPS. Nature 614, 334–342. 10.1038/s41586-022-05645-6.

19. LaMarche, N.M., Hegde, S., Park, M.D., Maier, B.B., Troncoso, L., Berichel, J.L., Hamon, P., Belabed, M., Mattiuz, R., Hennequin, C., et al. (2024). An IL-4 signalling axis in bone marrow drives pro-tumorigenic myelopoiesis. Nature 625, 166–174. 10.1038/s41586-023-06797-9.

20. Montfort, A., Colacios, C., Levade, T., Andrieu-Abadie, N., Meyer, N., and Ségui, B. (2019). The TNF Paradox in Cancer Progression and Immunotherapy. Front. Immunol. 10, 3666–5. 10.3389/fimmu.2019.01818.

21. Croft, M. (2009). The role of TNF superfamily members in T-cell function and diseases. Nat Rev Immunol 9, 271–285. 10.1038/nri2526.

22. Bankhead, P., Loughrey, M.B., Fernández, J.A., Dombrowski, Y., McArt, D.G., Dunne, P.D., McQuaid, S., Gray, R.T., Murray, L.J., Coleman, H.G., et al. (2017). QuPath: Open source software for digital pathology image analysis. Sci Rep-uk 7, 16878. 10.1038/s41598-017-17204-5.

23. Greenwald, N.F., Miller, G., Moen, E., Kong, A., Kagel, A., Dougherty, T., Fullaway, C.C., McIntosh, B.J., Leow, K.X., Schwartz, M.S., et al. (2021). Whole-cell segmentation of tissue images with human-level performance using large-scale data annotation and deep learning. Nat Biotech, 1–23. 10.1038/s41587-021-01094-0.

24. Backman, M., Strell, C., Lindberg, A., Mattsson, J.S.M., Elfving, H., Brunnström, H., O’Reilly, A., Bosic, M., Gulyas, M., Isaksson, J., et al. (2023). Spatial immunophenotyping of the tumour microenvironment in non–small cell lung cancer. Eur. J. Cancer 185, 40–52. 10.1016/j.ejca.2023.02.012.

25. Network, T.C.G.A.R. (2014). Comprehensive molecular profiling of lung adenocarcinoma. Nature 511, 1–9. 10.1038/nature13385.

26. Hammerman, P.S., Lawrence, M.S., Voet, D., Jing, R., Cibulskis, K., Sivachenko, A., Stojanov, P., McKenna, A., Lander, E.S., Gabriel, S., et al. (2012). Comprehensive genomic characterization of squamous cell lung cancers. Nature 489, 519–525. 10.1038/nature11404.

27. Datar, I.J., Hauc, S.C., Desai, S., Gianino, N., Henick, B., Liu, Y., Syrigos, K., Rimm, D.L., Kavathas, P., Ferrone, S., et al. (2021). Spatial Analysis and Clinical Significance of HLA Class-I and Class-II Subunit Expression in Non-Small Cell Lung Cancer. Clinical Cancer Research 27, 2837–2847. 10.1158/1078-0432.ccr-20-3655.

28. Montesion, M., Murugesan, K., Jin, D.X., Sharaf, R., Sanchez, N., Guria, A., Minker, M., Li, G., Fisher, V., Sokol, E.S., et al. (2021). Somatic HLA Class I Loss Is a Widespread Mechanism of Immune Evasion Which Refines the Use of Tumor Mutational Burden as a Biomarker of Checkpoint Inhibitor Response. Cancer Discov 11, 282–292. 10.1158/2159-8290.cd-20-0672.

29. Enfield, K.S.S., Colliver, E., Lee, C.S.Y., Magness, A., Moore, D.A., Sivakumar, M., Grigoriadis, K., Pich, O., Karasaki, T., Hobson, P.S., et al. (2024). Spatial Architecture of Myeloid and T Cells Orchestrates Immune Evasion and Clinical Outcome in Lung Cancer. Cancer Discov. 14, 1018–1047. 10.1158/2159-8290.cd-23-1380.

30. Burel, J.G., Pomaznoy, M., Arlehamn, C.S.L., Weiskopf, D., Antunes, R. da S., Jung, Y., Babor, M., Schulten, V., Seumois, G., Greenbaum, J.A., et al. (2019). Circulating T cell-monocyte complexes are markers of immune perturbations. eLife 8, e46045. 10.7554/elife.46045.

31. Burel, J.G., Pomaznoy, M., Arlehamn, C.S.L., Seumois, G., Vijayanand, P., Sette, A., and Peters, B. (2020). The Challenge of Distinguishing Cell–Cell Complexes from Singlet Cells in Non□Imaging Flow Cytometry and Single□Cell Sorting. Cytom. Part A 97, 1127–1135. 10.1002/cyto.a.24027.

32. Simoni, Y., Becht, E., Fehlings, M., Loh, C.Y., Koo, S.-L., Teng, K.W.W., Yeong, J.P.S., Nahar, R., Zhang, T., Kared, H., et al. (2018). Bystander CD8+ T cells are abundant and phenotypically distinct in human tumour infiltrates. Nature 557, 1–21. 10.1038/s41586-018-0130-2.

33. Kortekaas, K.E., Santegoets, S.J., Sturm, G., Ehsan, I., Egmond, S.L. van, Finotello, F., Trajanoski, Z., Welters, M.J.P., Poelgeest, M.I.E. van, and Burg, S.H. van der (2020). CD39 Identifies the CD4+ Tumor-Specific T-cell Population in Human Cancer. Cancer Immunol. Res. 8, 1311–1321. 10.1158/2326-6066.cir-20-0270.

34. Kim, J., Park, S., Kim, J., Kim, Y., Yoon, H.M., Rayhan, B.R., Jeong, J., Bothwell, A.L.M., and Shin, J.H. (2025). Trogocytosis-mediated immune evasion in the tumor microenvironment. Exp. Mol. Med. 57, 1–12. 10.1038/s12276-024-01364-2.

35. Steinert, E.M., Schenkel, J.M., Fraser, K.A., Beura, L.K., Manlove, L.S., Igyártó, B.Z., Southern, P.J., and Masopust, D. (2015). Quantifying Memory CD8 T Cells Reveals Regionalization of Immunosurveillance. Cell 161, 737–749. 10.1016/j.cell.2015.03.031.

36. Hedrick, C.C., and Malanchi, I. (2022). Neutrophils in cancer: heterogeneous and multifaceted. Nat. Rev. Immunol. 22, 173–187. 10.1038/s41577-021-00571-6.

37. Quail, D.F., Amulic, B., Aziz, M., Barnes, B.J., Eruslanov, E., Fridlender, Z.G., Goodridge, H.S., Granot, Z., Hidalgo, A., Huttenlocher, A., et al. (2022). Neutrophil phenotypes and functions in cancer: A consensus statement. J. Exp. Med. 219, e20220011. 10.1084/jem.20220011.

38. Wculek, S.K., and Malanchi, I. (2015). Neutrophils support lung colonization of metastasis-initiating breast cancer cells. Nature 528, 413–417. 10.1038/nature16140.

39. Saini, M., Szczerba, B.M., and Aceto, N. (2019). Circulating Tumor Cell-Neutrophil Tango along the Metastatic Process. Cancer Res. 79, 6067–6073. 10.1158/0008-5472.can-19-1972.

40. Szczerba, B.M., Castro-Giner, F., Vetter, M., Krol, I., Gkountela, S., Landin, J., Scheidmann, M.C., Donato, C., Scherrer, R., Singer, J., et al. (2019). Neutrophils escort circulating tumour cells to enable cell cycle progression. Nature 566, 553–557. 10.1038/s41586-019-0915-y.

41. Dickson, R.P., Erb-Downward, J.R., Martinez, F.J., and Huffnagle, G.B. (2015). The Microbiome and the Respiratory Tract. Annu. Rev. Physiol. 78, 1–24. 10.1146/annurev-physiol-021115-105238.

42. Hsu-Kim, C., Hoag, J.B., Cheng, G.-S., and Lund, M.E. (2013). The Microbiology of Postobstructive Pneumonia in Lung Cancer Patients. J. Bronchol. Interv. Pulmonol. 20, 266–270. 10.1097/lbr.0b013e31829ddf01.

43. Littman, A.J., Jackson, L.A., and Vaughan, T.L. (2005). Chlamydia pneumoniae and Lung Cancer: Epidemiologic Evidence. Cancer Epidemiology Prev. Biomark. 14, 773–778. 10.1158/1055-9965.epi-04-0599.

44. Jin, C., Lagoudas, G.K., Zhao, C., Bullman, S., Bhutkar, A., Hu, B., Ameh, S., Sandel, D., Liang, X.S., Mazzilli, S., et al. (2019). Commensal Microbiota Promote Lung Cancer Development via γδ T Cells. Cell 176, 998–1013.e16. 10.1016/j.cell.2018.12.040.

45. Tekle, G.E., and Garrett, W.S. (2023). Bacteria in cancer initiation, promotion and progression. Nat. Rev. Cancer 23, 600–618. 10.1038/s41568-023-00594-2.

46. Haastert, B., Mellanby, R.J., Anderton, S.M., and O’Connor, R.A. (2013). T Cells at the Site of Autoimmune Inflammation Show Increased Potential for Trogocytosis. PLoS ONE 8, e81404. 10.1371/journal.pone.0081404.

47. Ochs, J., Nissimov, N., Torke, S., Freier, M., Grondey, K., Koch, J., Klein, M., Feldmann, L., Gudd, C., Bopp, T., et al. (2022). Proinflammatory CD20+ T cells contribute to CNS-directed autoimmunity. Sci. Transl. Med. 14, eabi4632. 10.1126/scitranslmed.abi4632.

48. Hwang, I., Huang, J.-F., Kishimoto, H., Brunmark, A., Peterson, P.A., Jackson, M.R., Surh, C.D., Cai, Z., and Sprent, J. (2000). T Cells Can Use Either T Cell Receptor or Cd28 Receptors to Absorb and Internalize Cell Surface Molecules Derived from Antigen-Presenting Cells. J. Exp. Med. 191, 1137–1148. 10.1084/jem.191.7.1137.

49. Wetzel, S.A., McKeithan, T.W., and Parker, D.C. (2005). Peptide-Specific Intercellular Transfer of MHC Class II to CD4+ T Cells Directly from the Immunological Synapse upon Cellular Dissociation. J. Immunol. 174, 80–89. 10.4049/jimmunol.174.1.80.

50. Perez-Ruiz, E., Minute, L., Otano, I., Alvarez, M., Ochoa, M.C., Belsue, V., Andrea, C. de, Rodriguez-Ruiz, M.E., Perez-Gracia, J.L., Marquez-Rodas, I., et al. (2019). Prophylactic TNF blockade uncouples efficacy and toxicity in dual CTLA-4 and PD-1 immunotherapy. Nature 569, 428–432. 10.1038/s41586-019-1162-y.

51. Vasanthakumar, A., Liao, Y., Teh, P., Pascutti, M.F., Oja, A.E., Garnham, A.L., Gloury, R., Tempany, J.C., Sidwell, T., Cuadrado, E., et al. (2017). The TNF Receptor Superfamily-NF-κB Axis Is Critical to Maintain Effector Regulatory T Cells in Lymphoid and Non-lymphoid Tissues. Cell Reports 20, 2906–2920. 10.1016/j.celrep.2017.08.068.

52. Sharma, V.J., Starkey, G., D’Costa, R., James, F., Mouhtouris, E., Davis, L., Wang, B.Z., Vago, A., Raman, J., Mackay, L.K., et al. (2022). Australian Donation and Transplantation Biobank: A Research Biobank Integrated Within a Deceased Organ and Tissue Donation Program. Transplant Direct 9, e1422. 10.1097/txd.0000000000001422.

53. Preibisch, S., Saalfeld, S., and Tomancak, P. (2009). Globally optimal stitching of tiled 3D microscopic image acquisitions. Bioinformatics 25, 1463–1465. 10.1093/bioinformatics/btp184.

54. Valen, D.A.V., Kudo, T., Lane, K.M., Macklin, D.N., Quach, N.T., DeFelice, M.M., Maayan, I., Tanouchi, Y., Ashley, E.A., and Covert, M.W. (2016). Deep Learning Automates the Quantitative Analysis of Individual Cells in Live-Cell Imaging Experiments. PLoS Comput. Biol. 12, e1005177. 10.1371/journal.pcbi.1005177.

55. Windhager, J., Zanotelli, V.R.T., Schulz, D., Meyer, L., Daniel, M., Bodenmiller, B., and Eling, N. (2023). An end-to-end workflow for multiplexed image processing and analysis. Nat. Protoc. 18, 3565–3613. 10.1038/s41596-023-00881-0.

56. Pau, G., Fuchs, F., Sklyar, O., Boutros, M., and Huber, W. (2010). EBImage—an R package for image processing with applications to cellular phenotypes. Bioinformatics 26, 979–981. 10.1093/bioinformatics/btq046.

57. McLachlan (2005). Discriminant analysis and statistical pattern recognition (John Wiley & Sons).

58. Phipson, B., Sim, C.B., Porrello, E.R., Hewitt, A.W., Powell, J., and Oshlack, A. (2022). propeller: testing for differences in cell type proportions in single cell data. Bioinformatics 38, 4720–4726. 10.1093/bioinformatics/btac582.

59. Daubeuf, S., Puaux, A.-L., Joly, E., and Hudrisier, D. (2006). A simple trogocytosis-based method to detect, quantify, characterize and purify antigen-specific live lymphocytes by flow cytometry, via their capture of membrane fragments from antigen-presenting cells. Nat. Protoc. 1, 2536–2542. 10.1038/nprot.2006.400.

60. Sanchez-Cespedes, M., Parrella, P., Esteller, M., Nomoto, S., Trink, B., Engles, J.M., Westra, W.H., Herman, J.G., and Sidransky, D. (2002). Inactivation of LKB1/STK11 is a common event in adenocarcinomas of the lung. Cancer Res. 62, 3659–3662.

61. Liu, N., Bhuva, D.D., Mohamed, A., Bokelund, M., Kulasinghe, A., Tan, C.W., and Davis, M.J. (2023). standR: spatial transcriptomic analysis for GeoMx DSP data. Nucleic Acids Res. 52, e2–e2. 10.1093/nar/gkad1026.

62. Robinson, M.D., and Oshlack, A. (2010). A scaling normalization method for differential expression analysis of RNA-seq data. Genome Biol. 11, R25. 10.1186/gb-2010-11-3-r25.

63. Law, C.W., Chen, Y., Shi, W., and Smyth, G.K. (2014). voom: Precision weights unlock linear model analysis tools for RNA-seq read counts. Genome Biol. 15, R29. 10.1186/gb-2014-15-2-r29.

64. Robinson, M.D., McCarthy, D.J., and Smyth, G.K. (2009). edgeR: a Bioconductor package for differential expression analysis of digital gene expression data. Bioinformatics 26, 139–140. 10.1093/bioinformatics/btp616.

65. Smyth, G.K. (2011). Linear Models and Empirical Bayes Methods for Assessing Differential Expression in Microarray Experiments. Statistical Applications in Genetics and Molecular Biology 3, 1–25. 10.2202/1544-6115.1027.

66. Phipson, B., Lee, S., Majewski, I.J., Alexander, W.S., and Smyth, G.K. (2016). Robust hyperparameter estimation protects against hypervariable genes and improves power to detect differential expression. Ann. Appl. Stat. 10, 946–963. 10.1214/16-aoas920.

67. Y, B.Y.H. (1995). Controlling_The_False_Discovery_Rate_-_A_Practical. statistical 57, 289–300. 10.2307/2346101.

68. McCarthy, D.J., and Smyth, G.K. (2009). Testing significance relative to a fold-change threshold is a TREAT. Bioinformatics 25, 765–771. 10.1093/bioinformatics/btp053.

69. Liberzon, A., Birger, C., Thorvaldsdóttir, H., Ghandi, M., Mesirov, J.P., and Tamayo, P. (2015). The Molecular Signatures Database Hallmark Gene Set Collection. Cell Syst. 1, 417–425. 10.1016/j.cels.2015.12.004.

70. Wu, D., and Smyth, G.K. (2012). Camera: a competitive gene set test accounting for inter-gene correlation. Nucleic Acids Res. 40, e133–e133. 10.1093/nar/gks461.

71. Ritchie, M.E., Phipson, B., Wu, D., Hu, Y., Law, C.W., Shi, W., and Smyth, G.K. (2015). limma powers differential expression analyses for RNA-sequencing and microarray studies. Nucleic Acids Res 43, e47–e47. 10.1093/nar/gkv007.

