## Supplementary figures and images for "TNF⍰-mediated myeloid-instructed CD14^+^CD4^+^ T cells within the tumor microenvironment are associated with poor survival in non-small cell lung cancer"

### Supp Figures

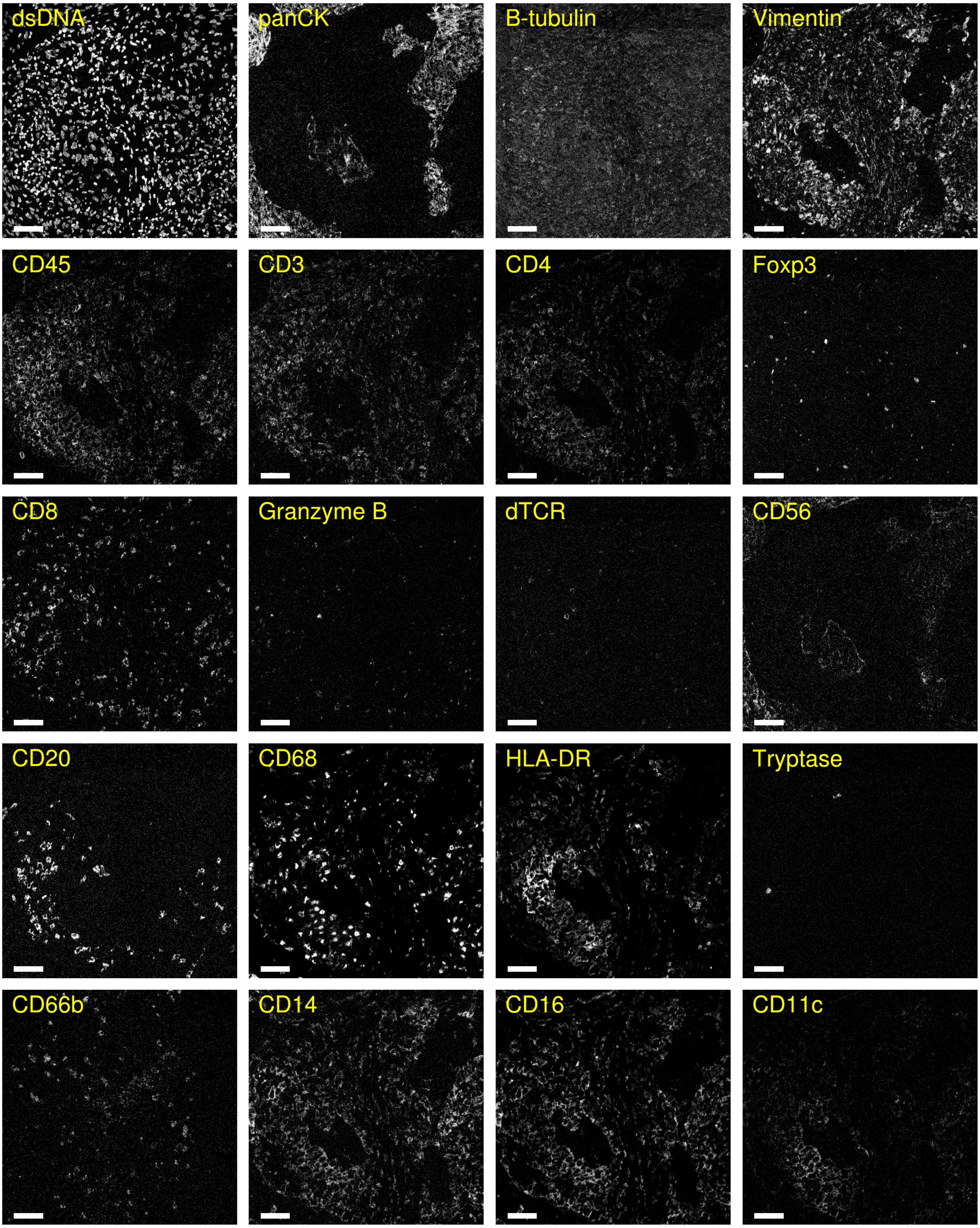

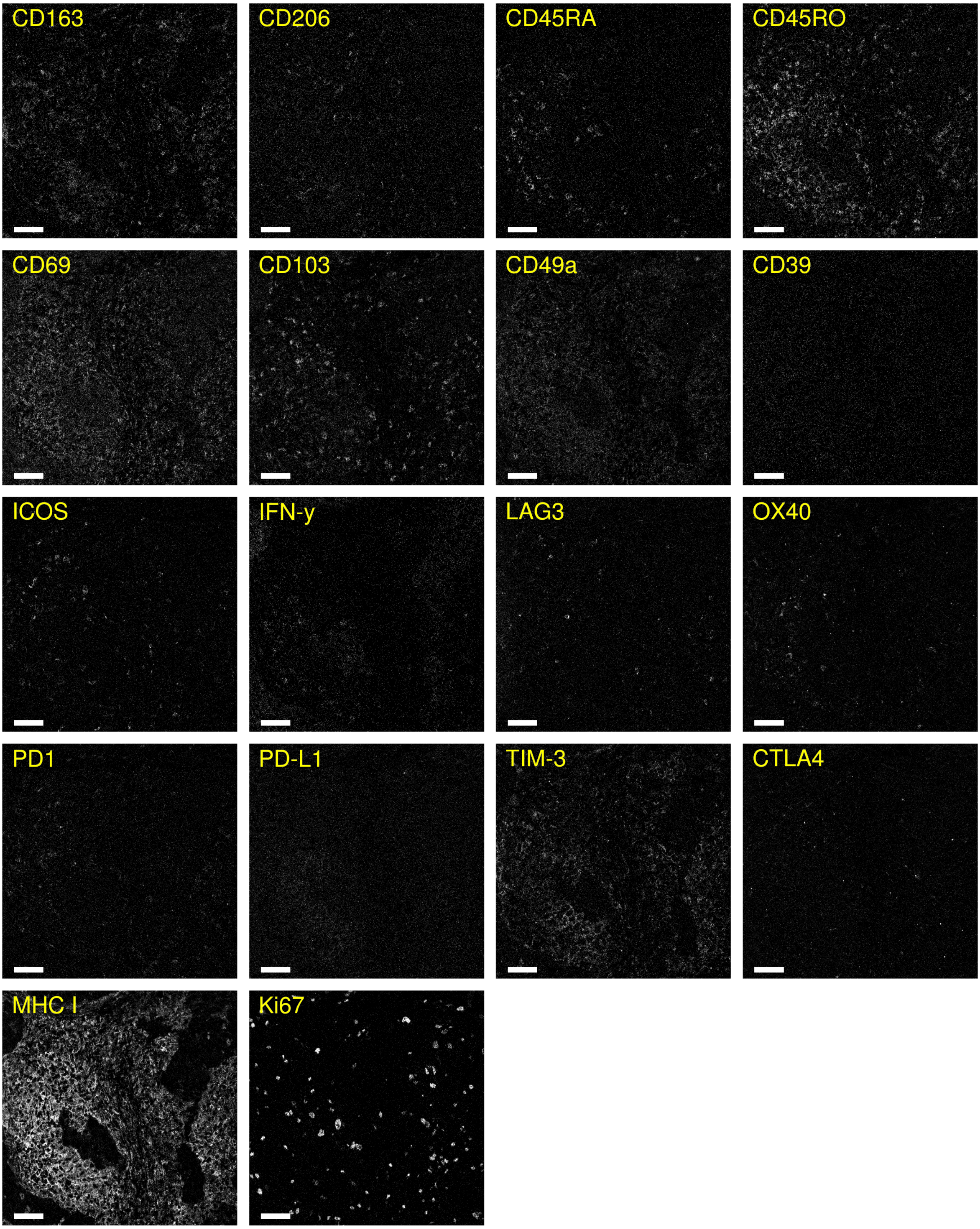

B

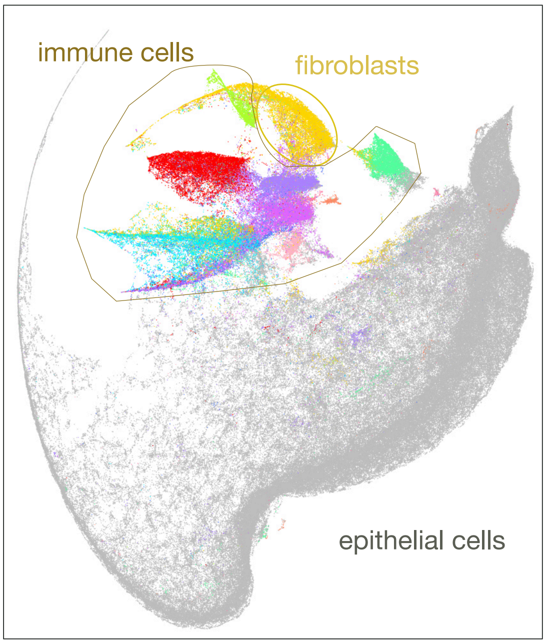

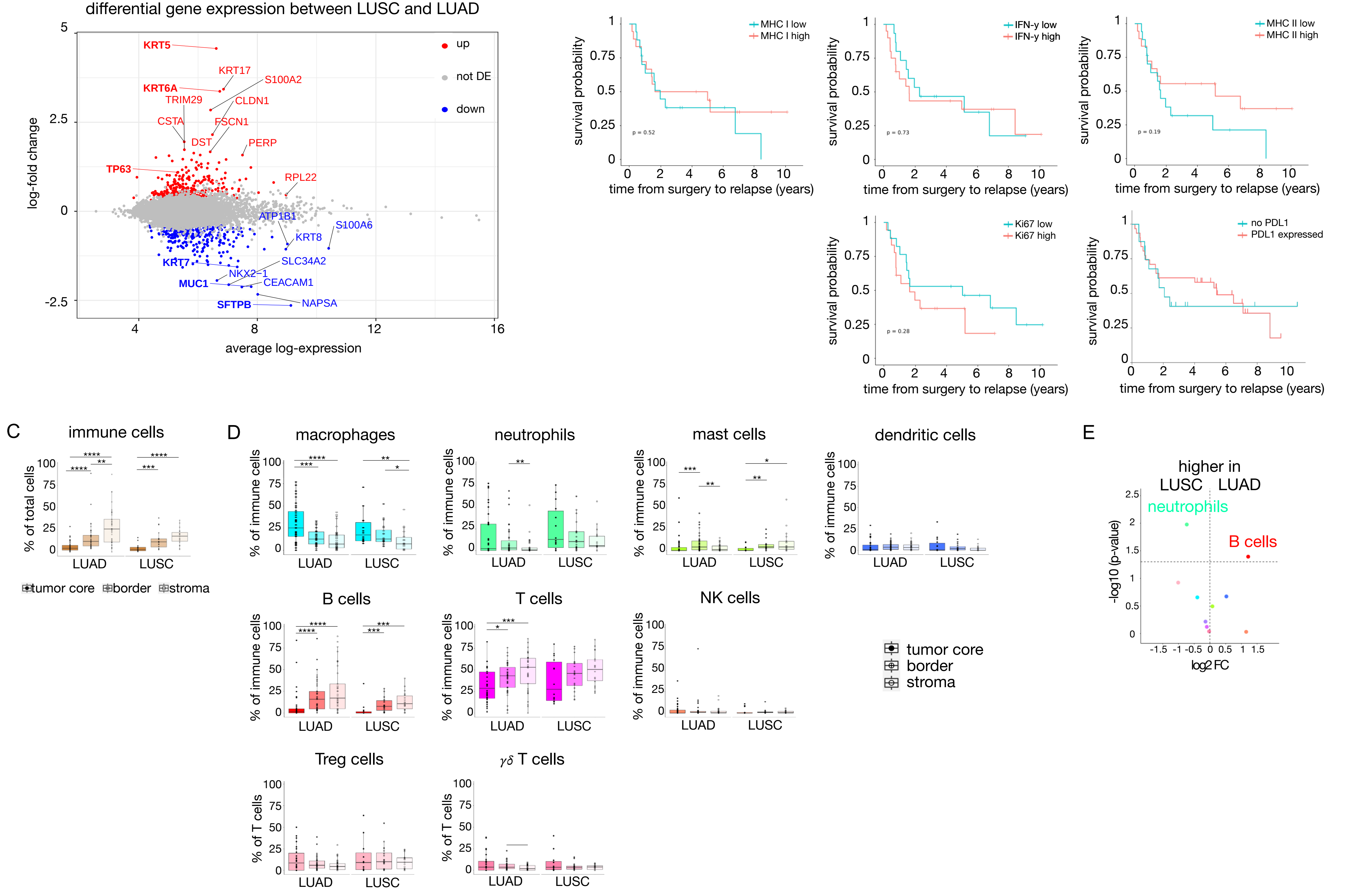

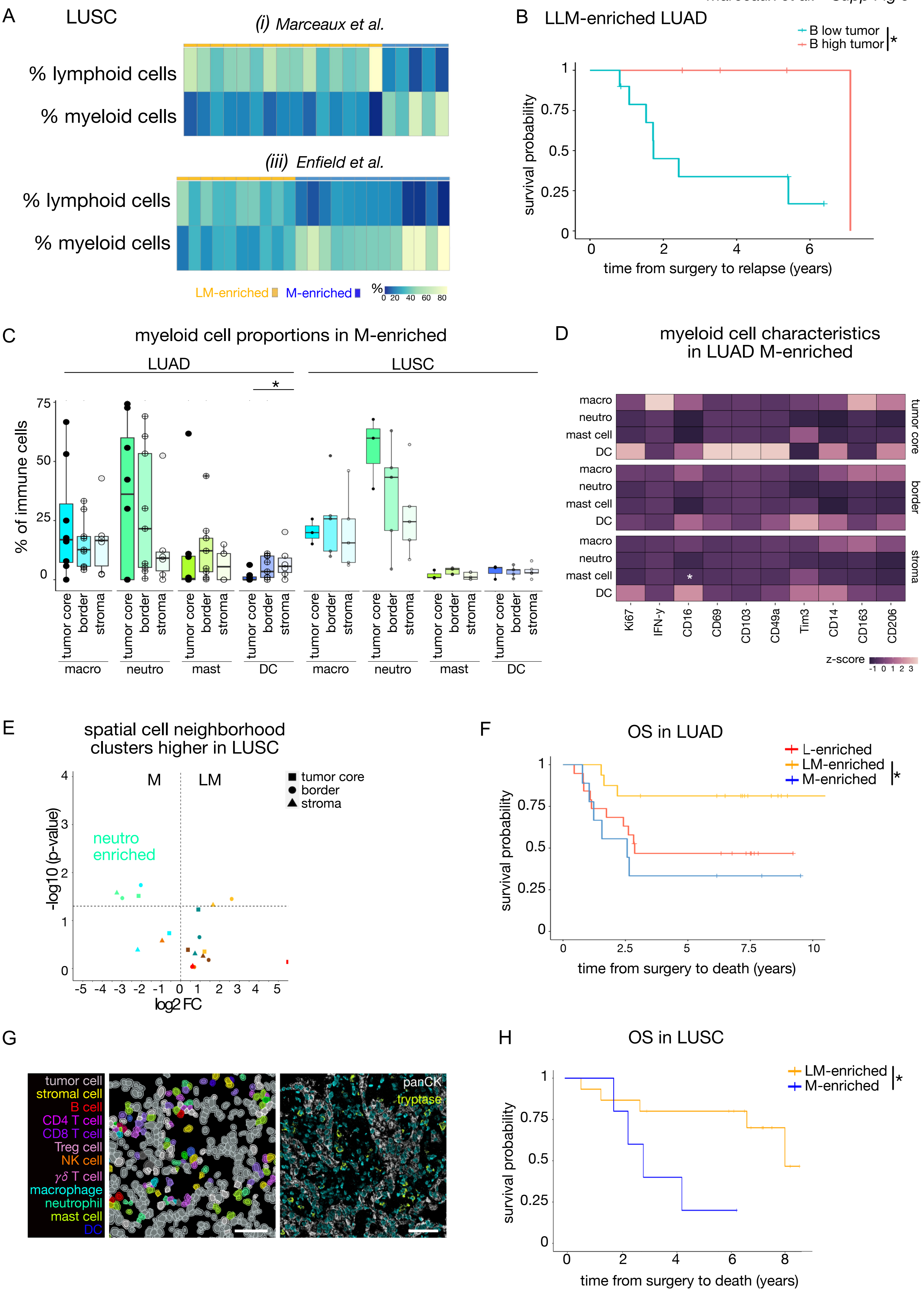

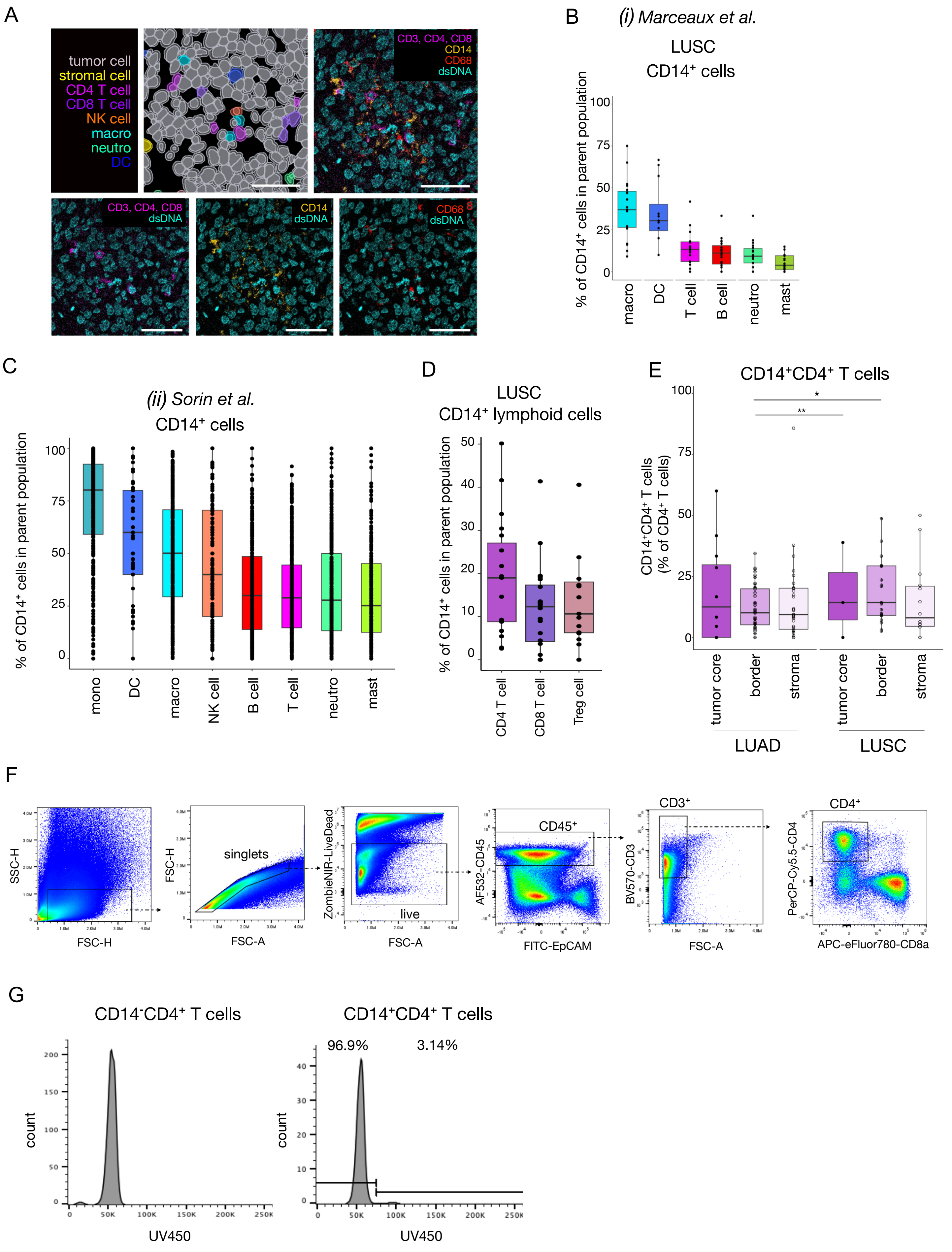

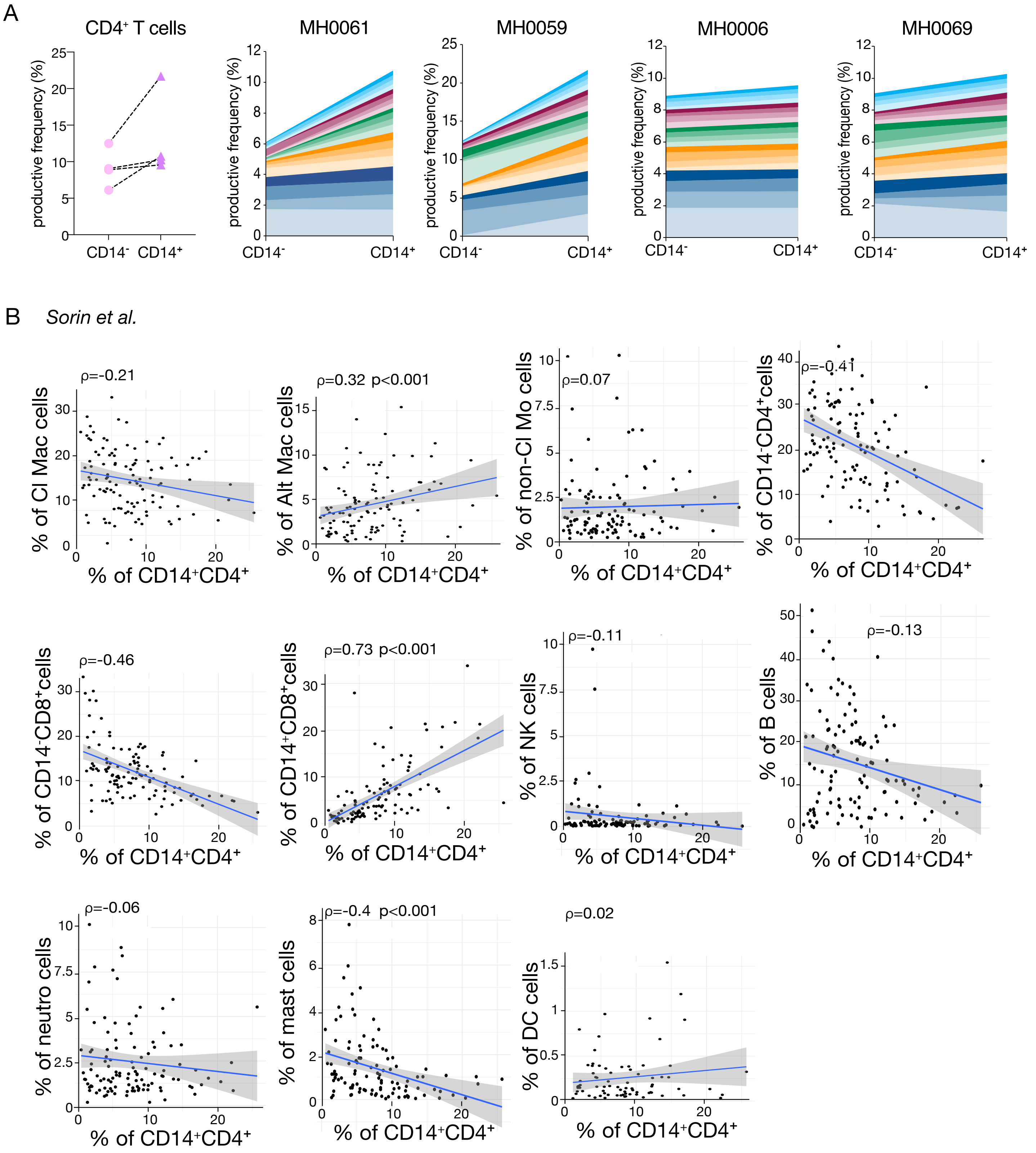

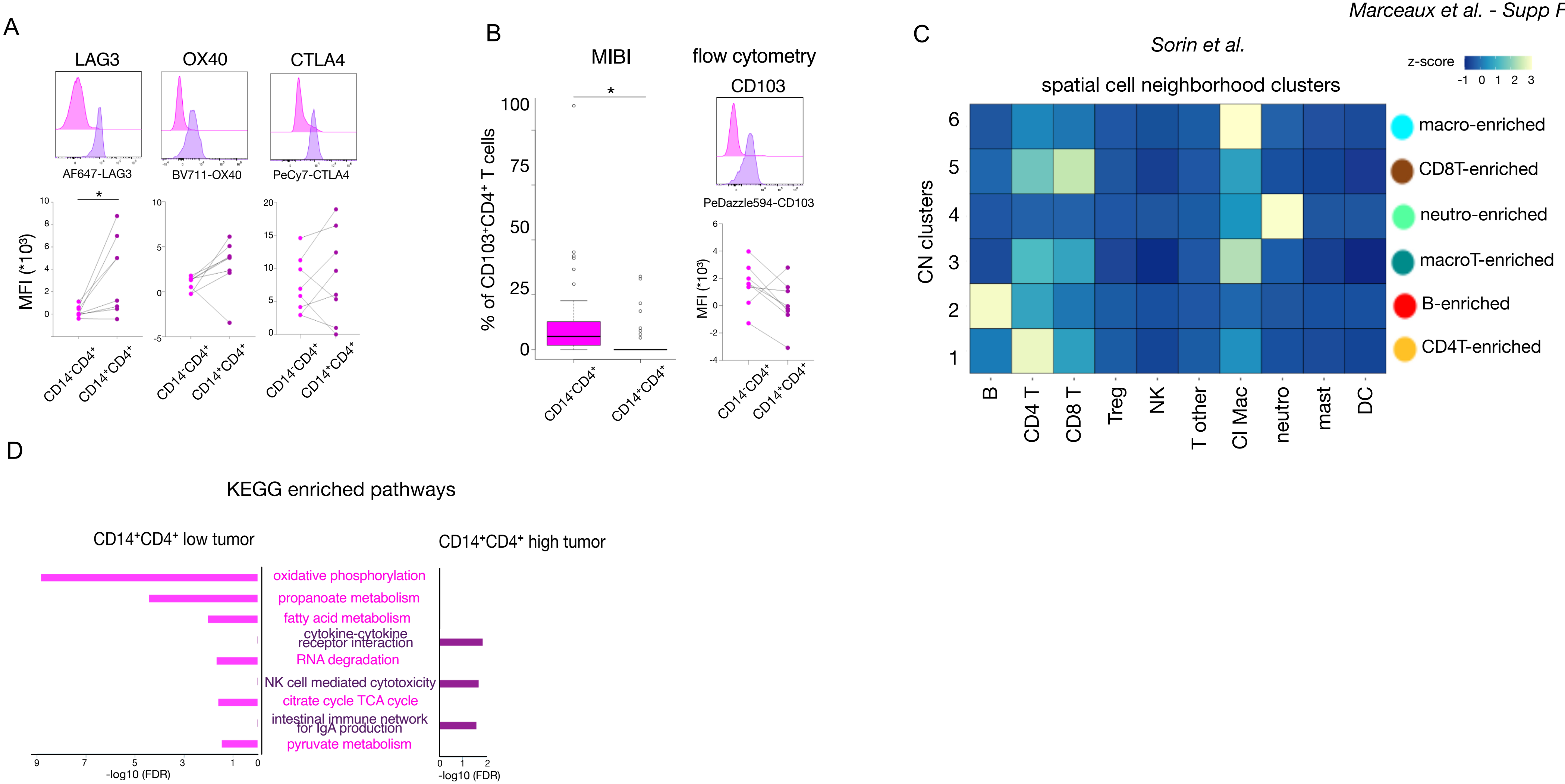
